# Cryo-EM structure of the *Mycobacterium abscessus* bacteriophage ФJabs

**DOI:** 10.64898/2026.07.24.740622

**Authors:** Christian Cambillau, Jun Hao Liew, Bernice Siu Yan Tan, Laurent Kremer, Pablo Bifani, Adeline Goulet

## Abstract

Exploring bacteriophage structural diversity is crucial for advancing our understanding of phage biology and therapeutic applications. Yet, high-resolution structures of phages infecting mycobacteria remain scarce. Here, we present the cryo-electron microscopy structure of ФJabs, a siphophage infecting the multidrug-resistant human pathogen *Mycobacterium abscessus,* which combines several unusual architectural features. Its T=9 icosahedral capsid is assembled from two distinct major capsid proteins, one forming the hexons and the other the pentons, a feature previously assigned to prolate capsid only. An extensive network of ∼2,000 disulfide bonds stabilizes individual structural components and establishes covalent links between the connector, tail and baseplate. The distal tail features a complex baseplate comprising multiple candidate receptor-binding proteins containing carbohydrate-binding modules and β-sandwich domains, the latter organized into uncommon heterotrimeric assemblies. These findings expand our understanding of phage structural diversity and provide structural insights for future investigations of the molecular determinants of host specificity.

## INTRODUCTION

The nearly 16,000 bacteriophages (phages) known to infect mycobacteria (mycobacteriophages; https://phagesdb.org/) constitute a vast yet largely unexplored resource for understanding phage structural and functional diversity ^1,2^. Beyond their fundamental interest, mycobacteriophages are emerging as promising therapeutic agents against multidrug-resistant human pathogens ^3,4,5,6^. Despite their remarkable diversity and growing clinical relevance, high-resolution information on mycobacteriophage virions remains limited ^3,7–10,11,12^, restricting our understanding of the structural principles underlying host recognition and infection.

This limitation is especially significant for phages infecting *Mycobacterium abscessus* (*Mabs*), an emerging multidrug-resistant opportunistic pathogen responsible for severe pulmonary disease ^13^. The therapeutic potential of mycobacteriophages was demonstrated through the compassionate use of a three-mycobacteriophage cocktail that successfully treated a cystic fibrosis patient with a disseminated *Mabs* infection ^14^. To date, no high-resolution structure of *Mabs*-infecting phage is available. We previously predicted the architectures of baseplates from various mycobacteriophages, revealing these tail tip multiprotein assemblies to be highly modular and structurally diverse, with domains predicted to recognize sugar- and lipid-containing components of the mycobacterial cell envelope ^15^. However, how these proteins assemble into functional baseplates and are integrated into complete infectious virions remains poorly understood.

This question is particularly relevant for ФJabs, a siphophage infecting *Mabs* ^4^. During the course of *Mabs* infection, the bacterium can transition from a smooth (S) to a rough (R) variant, mainly through the loss of surface glycopeptidolipids (GPL), resulting in enhanced virulence and immune evasion ^16,17–19^. While all previously characterized *Mabs* phages preferentially infect the GPL-deficient R morphotype ^14,20^, ФJabs is the first phage reported to preferentially infect the GPL-rich S morphotype ^4,14^. Conversely, the S-to-R transition, resulting from mutations in genes coding for GPL biosynthetic enzymes or transport, confers resistance to ФJabs, suggesting recognition of a distinct receptor, potentially involving GPL ^4^. Understanding the structural and molecular basis of ФJabs host specificity is, therefore, important for uncovering principles governing phage-host interactions and for designing therapeutic phage cocktails targeting both *Mabs* morphotypes.

Here, we determined the cryo-electron microscopy (cryo-EM) structure of ФJabs and built atomic models of its major virion components. Although ФJabs displays the canonical siphophage organization, it combines several unusual architectural features, including a T=9 icosahedral capsid assembled from two distinct major capsid proteins, a simplified three-component connector lacking an identifiable stopper protein, an elaborate baseplate containing multiple candidate receptor-binding proteins including a heterotrimeric assembly of β-sandwich domains, and an extensive covalent network of disulfide bonds that stabilizes individual components of the virion and links the connector, tail and baseplate. Together, these findings expand our understanding of mycobacteriophage structural diversity and establish a structural framework for future studies of *Mabs* variant specificity.

## RESULTS

We determined the cryo-EM structure of the ФJabs virion and built atomic models for its major components (Fig. 1a,b, and Supplementary Fig. S1 to S4, and Table S1). To support accurate structural modelling of ФJabs, we performed additional Nanopore sequencing to resolve split-gene annotations affecting the portal protein, the tape measure protein (TMP), and a protein identified structurally as a spike protein (Supplementary Table S2 and Fig. 1c). The sequence of another virion protein, which we designated β-sandwich 2 based on its overall fold (Fig. 1b,c), was also revised (H46 versus P46; Supplementary Table S2). The revised genome annotation comprises 15 genes encoding virion proteins (Fig. 1c). The ФJabs virion consists of an icosahedral capsid linked through a head-tail connector to a long, flexible, non-contractile tail that terminates in a distal baseplate (Fig. 1a). We determined the structures of the major virion components, the capsid, connector, tail tube, and baseplate, and thereby assembled a model the complete phage particle (Fig. 1b, and Supplementary Fig. S1 and S2, and Table S1).

**Fig. 1.**
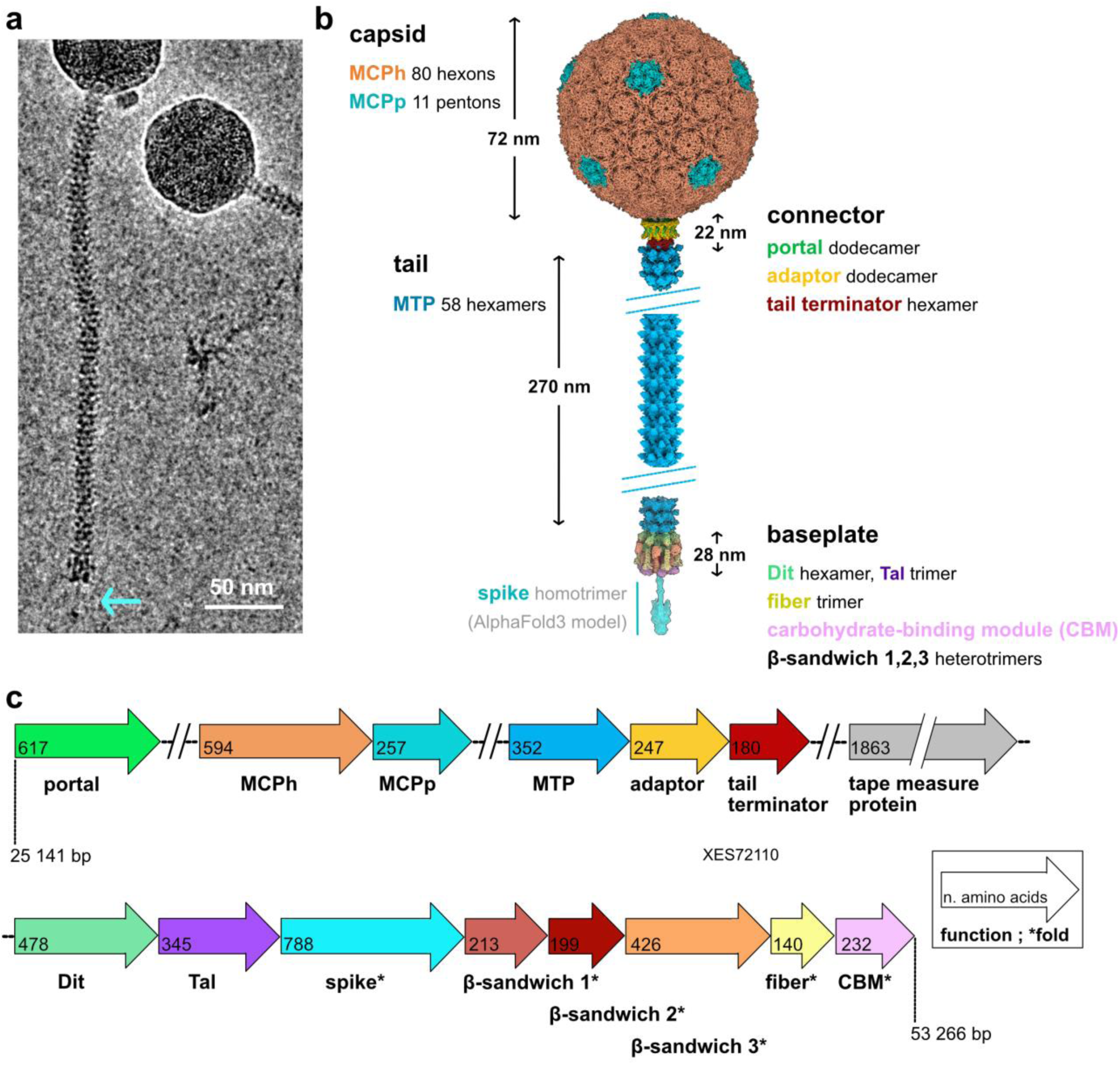
ФJabs cryo-EM structure. **a.** Cryo-EM micrograph of individual virions. **b.** 3D reconstructions of the ФJabs capsid, connector, baseplate, and tail segments bound to the connector and baseplate. The long tail segment between the white lines is shown as a surface rendering of juxtaposed MTP structural models to illustrate the tail. The AlphaFold3-predicted structure of spike trimer is shown as ribbon and surface representations to illustrate the complete virion. The corresponding arrow-like spike is visible in the micrograph in **a** (indicated with a blue arrow). **c** Schematic representation of the genomic region spanning from the *portal* to the *cbm* genes (color code as in b). The inset explains the information provided within and below each arrow, corresponding to the number of amino acids and the functional annotation, respectively. The asterisk (*) indicates virion proteins named according to their structural features.

### Two major capsid proteins assemble a disulfide-cross-linked T=9 icosahedral capsid

We reconstructed the ФJabs capsid with imposed icosahedral symmetry to an overall resolution of 3.4 Å (Fig. 2a, and Supplementary Fig. S1 and S3, and Table S1). The 72-nm-diameter particle adopts a T=9 icosahedral geometry and encloses multiple concentric layers of dsDNA (Fig. 2a, and Supplementary Fig. S5a). The capsid shell is exclusively built from two major capsid proteins (MCP; Fig. 2a), with no additional cement or decoration proteins: MCPh (594 residues; Fig. 1c and table S2) forms the 80 hexons, whereas MCPp (257 residues; Fig. 1c and Supplementary Table S2) forms the 11 pentons, giving a total of 535 MCP subunits.

**Fig. 2.**
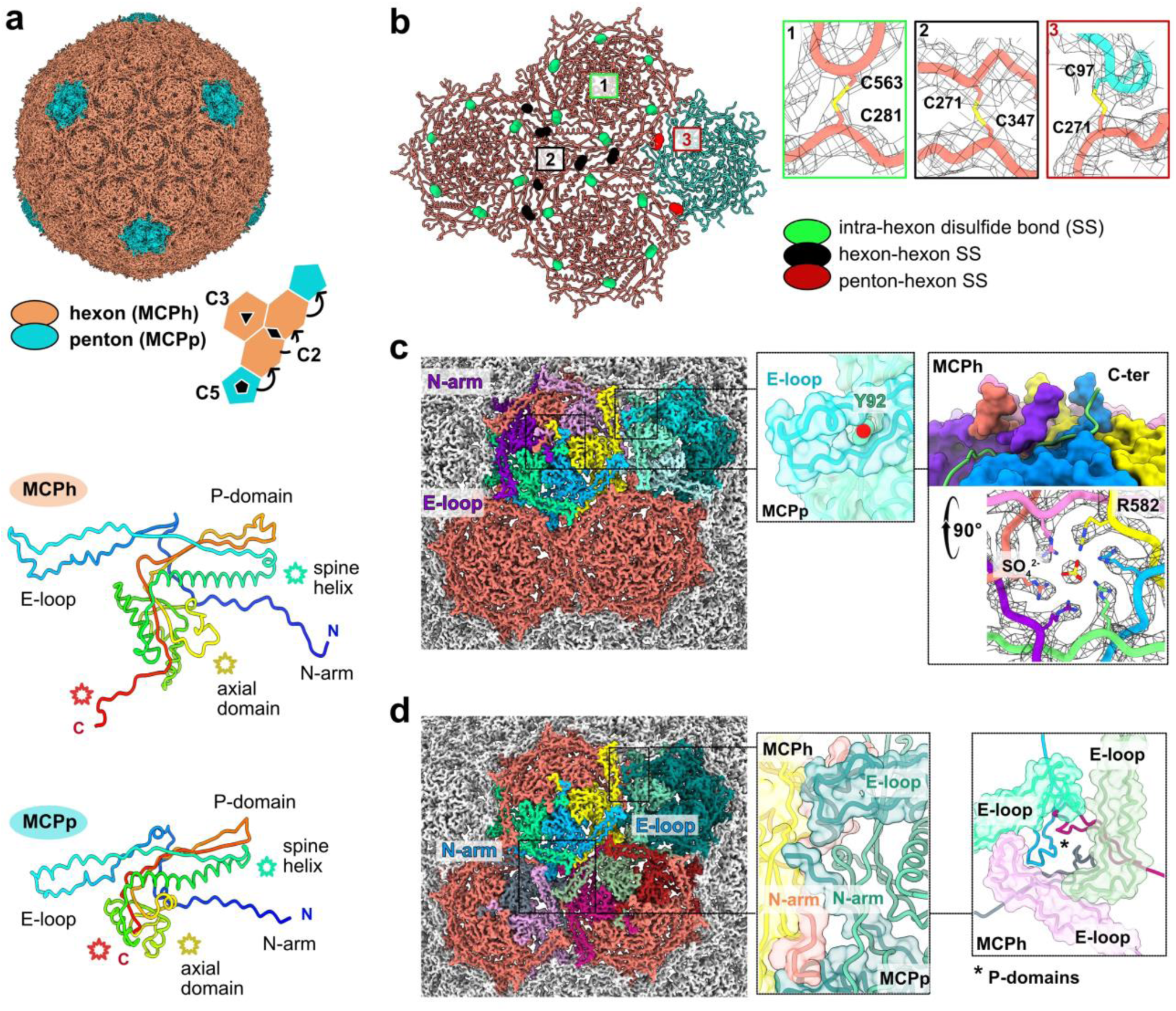
Two major capsid proteins assemble a disulfide-cross-linked T=9 icosahedral capsid. **a.** (top) 3D reconstruction of the capsid with imposed icosahedral symmetry. Hexons (MCPh) and pentons (MCPp) are colored salmon and turquoise, respectively. The schematic highlights the T=9 icosahedral geometry. (middle, bottom) Rainbow-colored ribbon representations of MCPh and MCPp. Star symbols indicate regions of structural divergence between the two proteins. **b.** (left) Ribbon representation of three hexons and one penton. Disulfide bonds within hexons (1) are shown as green spheres, whereas inter-capsomer disulfide bonds linking neighboring hexons (2) or hexons and pentons (3) are shown as black and red spheres, respectively. (right) Close-up views of the disulfide bonds shown as sticks with cysteine residues labeled. Cryo-EM density is shown as mesh. **c.** (left) Reconstruction with MCPh and MCPp subunits highlighted to illustrate intra-capsomer interactions. (middle) Close-up of the MCPp "knob-in-hole" interaction. The cryo-EM density is shown as surface and the residue Y92 as sphere. (right) Turret-like structure formed by the MCPh C-terminal extensions within a hexon. One MCPh subunit is shown as a ribbon for clarity, the others as surfaces. The density is shown as mesh and the assigned sulfate ion as sticks. **d.** (left) Reconstruction highlighting inter-capsomer interactions. (middle) N-arm and E-loop interacting with MCPh subunits of a neighboring hexon. (right) Junction of three hexons showing E-loops as ribbons and surfaces and the central P-domains as ribbons

Despite limited sequence identity (18% across the conserved region; alignment spanning residues 252-594 of the longer MCPh; fig. S5B), MCPh and MCPp both adopt the canonical HK97 fold ^21,22^, comprising an N-terminal arm (N-arm), an extended loop (E-loop), a spine α-helix, a peripheral domain (P-domain) and an axial domain (Fig. 2a, and Supplementary Fig. S5c). Structural similarity searches with Foldseek ^23^ returned significant matches to MCPs of mycobacteriophages such as Che8 ^7^, as well as other HK97-fold phages such as the *Escherichia coli* phage T5 (Supplementary Table S3 and Fig. S5d). We could model MCPp almost completely (residues 3-257), whereas density for MCPh residues 1-266 was absent and only residues 267-589 could be built. This observation strongly suggests proteolytic removal of an N-terminal maturation domain, consistent with the maturation pathway reported for HK97-like phages ^21,22^ and the presence of a gene annotated as encoding a capsid maturation protease (Genbank accession number XES72123).

Although they share the same overall fold (Supplementary Fig. S5b), MCPh and MCPp differ in several notable structural features (Fig. 2a (middle, bottom)). First, the spine helix of MCPp has a ∼25° bend that produces a distinct elbow-like conformation. Second, the axial domain of MCPp is markedly smaller than that of MCPh, owing to shorter loops and the absence of an α-helix. Third, MCPh carries a long, 17-residue C-terminal extension. Further differences in the conformations of the P-domain and the distal E-loop (Fig. 2a (middle, bottom)) reflect structural adaptations to mediate intra- and inter-capsomer interactions.

The N-arm, E-loop, and P-domain of MCPh and MCPp form an extensive interaction network connecting neighboring subunits and capsomers (Fig. 2b,c,d). Remarkably, each hexon is stabilized by six inter-subunit disulfide bonds, resulting in a total of 480 bonds across the 80 hexons, each linking C281 in the N-arm of one MCPh subunit to C563 in the linker between the P-domain and axial-domain of the adjacent MCPh subunit (Fig. 2b, and Supplementary Table S4). The hexon-hexon and penton-hexon interfaces are likewise reinforced by disulfide cross-links: two disulfide bonds per hexon-hexon interface form between MCPh residues C271 (N-arm) and C347 (E-loop), resulting in a total of 420 bonds across the 80 hexons, whereas one disulfide bond per penton-hexon interface, resulting in a total of 55 bonds across the 11 pentons, links MCPp C97 (P-domain) to MCPh C271 (N-arm) (Fig. 2b, and Supplementary Table S4). In total, the ФJabs capsid is stabilized by an extensive covalent framework of 955 disulfide bonds (Supplementary Table S4), further reinforced by numerous non-covalent contacts.

Within a capsomer, the N-arm of MCP subunit i primarily contacts the axial and P-domains of MCP subunit i-1, whereas its E-loop interacts with the spine helix and P-domain of MCP subunit i+1 (Fig. 2c). In pentons, the bulky MCPp residue Y92 forms a “knob-in-hole” interaction with the E-loop of the adjacent MCPp subunit. In hexons, the MCPh C-terminal extensions make long-range contacts with three adjacent MCP subunits (i-1, i+1, and i+2), and collectively, the six extensions assemble into a turret-like structure protruding from the center of each hexon (Fig. 2c). A well-defined density is observed at the center of this turret. Based on its coordination by the six MCPh R582 side chains, we tentatively assigned this density to a sulfate ion (SO4²̅), which may contribute to stabilizing the MCPh hexamer (Fig. 2c). However, the identity of this ligand cannot be unambiguously established. The resolution of our reconstruction does not allow us to reliably distinguish SO4²̅ from a chloride ion, as both are present in the phage buffer. Nevertheless, the refinement statistics are more consistent with the assignment of a sulfate ion.

Inter-capsomer contacts are mainly mediated by the MCP N-arms and E-loops (Fig. 2d). At both the penton-hexon and hexon-hexon interfaces, the N-arm contacts MCP subunits in the neighboring capsomer. In addition, the bent tip of the MCPp E-loop inserts into a shallow groove between two adjacent MCPh subunits of a neighboring hexon (Fig. 2d), thereby strengthening penton-hexon connectivity. Furthermore, at the junction of three hexons, the P-domains and E-loops of six MCPh subunits form a triangular arrangement centered on a local threefold axis (Fig. 2d). A similar arrangement occurs at the junction of two hexons and one penton, although the local symmetry is partly disrupted by conformational differences in the E-loop at these mixed MCPh-MCPp interfaces (Supplementary Fig. S5e).

### The portal, adaptor and tail terminator proteins form the connector

The connector occupies a unique fivefold vertex, replacing a penton and linking the capsid to the tail (Fig. 1a,b, and Fig. 3a). We reconstructed it without imposed symmetry to an overall resolution of 6 Å (Supplementary Fig. S1, S3, S6a, and Table S1). The ФJabs connector comprises a dodecameric portal protein embedded within the capsid, followed sequentially by a dodecameric adaptor and a hexameric tail terminator (TT), the last of which contacts the first hexameric ring of the major tail protein (MTP) (Fig. 3a). Notably, the connector lacks an identifiable stopper protein, a component commonly found in siphophage connector assemblies ^24,25^. Localized reconstructions of the portal-adaptor and TT-MTP assemblies, refined with C12 and C6 symmetry imposed, respectively, yielded maps at 3.4 Å and 3.9 Å resolution for model building (Fig. 3b,c,d, and Supplementary Fig. S1, S3, and Table S1). We then assembled and refined the resulting models within a C12-C6 composite map to generate the final connector-MTP structure (Fig. 3e, and Supplementary Fig. S1 and Table S1).

**Fig. 3.**
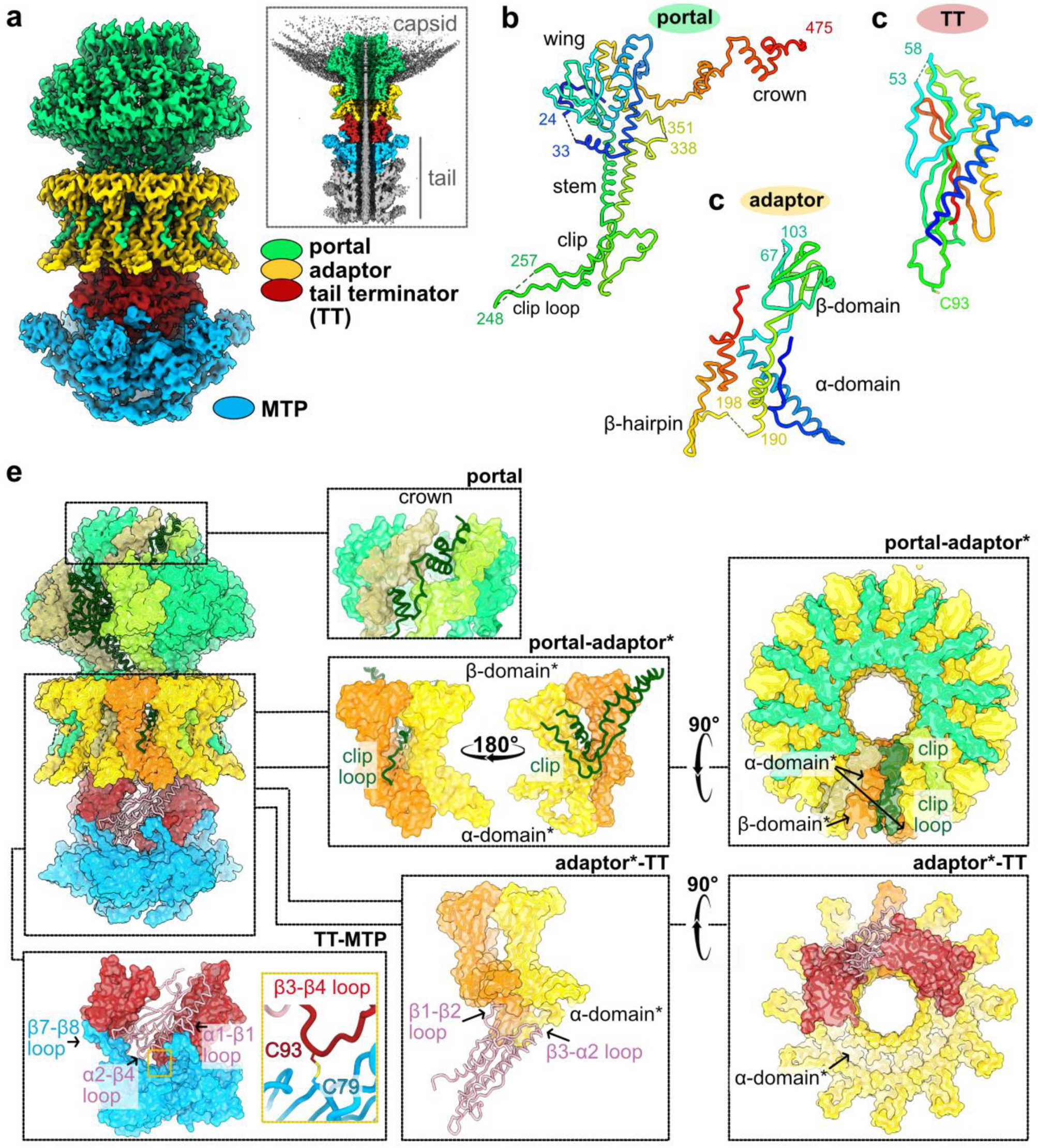
The portal, adaptor and tail terminator proteins form the connector. **a** Composite map of the connector bound to the first MTP ring, assembled from the C12-symmetrized portal-adaptor and C6-symmetrized TT-MTP localized reconstructions. Inset: cut-away view of the complete reconstruction highlighting the insertion of the connector into the capsid. **b-d.** Ribbon representations of the portal, adaptor and TT structures, respectively. **e.** Ribbon and surface representations of the connector-MTP assembly. Insets show close-up views of protein-protein interactions between portal subunits and at the portal-adaptor, adaptor-TT and TT-MTP interfaces.

We modeled 428 of the 617 residues of the ФJabs portal protein (Fig. 1c and Supplementary Table S2) into the cryo-EM density. The N-terminal (residues 1-11) and C-terminal (residues 475-617) regions, together with three internal loops (residues 25-32, 132-136, 249-256, and 337-351), lacked interpretable density. The portal protein adopts the canonical fold of tailed phage portal proteins, comprising crown, wing, stem, and clip domains (Fig. 3b, and Supplementary Table S3 and Fig. S6b). Two features nonetheless distinguish it from previously characterized portal proteins. First, the resolved part of the crown domain (residues 400-475) is unusually extended and contacts adjacent portal subunits, creating a continuous inter-subunit network that forms an interlocked ring at the top of the dodecamer (Fig. 3e). Second, the clip domain is substantially enlarged, though less well resolved than the portal central core. In particular, a prominent loop within the clip domain (residues 234-264), not fully modeled owing to weak density at its distal tip, projects outward from the portal channel and intercalates between two adjacent adaptor subunits, thereby contributing to portal-adaptor association (Fig. 3a,b,e).

We modeled residues 3-247 of the adaptor protein (247 residues; Fig. 1c and Supplementary Table S2), with the exception of residues 68-102 and 191-197, which lacked interpretable density (Fig. 3c). It comprises two domains: a predominantly α-helical domain formed by the N- and C-terminal regions (α-domain) and a β-stranded domain (β-domain, residues 59-166). The ФJabs adaptor protein is most similar to that of coliphage DT57C ^41^ (Supplementary Table S3 and Fig. S6c). The portal-adaptor interface is intricate: the β-domain and C-terminal extension of each adaptor subunit pack against the clip domain of one portal subunit at the periphery of the portal ring, generating a ring-within-a-ring architecture, while the clip loop of the portal protein is enclosed by the β- and α-domains of two adjacent adaptor subunits (Fig. 3e). The low-resolution asymmetric capsid-connector reconstruction additionally revealed the adaptor-capsid interface, showing interactions between five and seven non-consecutive MCPh and adaptor subunits, respectively, reflecting the C5-C12 symmetry mismatch (Supplementary Fig. S6a,d,e). Of the five interacting MCPh subunits, three contact the loops at the top of the β-domains of individual adaptor subunits through their E-loops (Supplementary Fig. S6f), whereas the remaining two engage the interfaces between adjacent adaptor subunits (Supplementary Fig. S6g).

Distal to the adaptor dodecamer, the TT hexamer links the connector to the tail tube (Fig. 3a). We modeled nearly the entire TT protein (180 residues; Fig. 1C and Supplementary Table S2), except for a disordered loop (residues 54-58). Its structure comprises two α-helices packed against a five-stranded β-sheet (Fig. 3d), a fold conserved among tailed phages and particularly similar to that of E. *coli* phage lambda ^26^ (Supplementary Table S3 and Fig. S6f). Each TT subunit inserts its β1-β2 and β3-α2 loops into the cleft formed by the α-domains of two adjacent adaptor subunits, thereby accommodating the C12-C6 symmetry mismatch at the adaptor-TT interface and anchoring the hexameric TT ring within the clamp-like base of the adaptor dodecamer (Fig. 3e). At its distal end, each TT subunit contacts two adjacent MTP subunits through its N-terminal helix and its α1-β1, β2-β3 and α2-β4 loops, which engage the prominent β7-β8 loop of the first MTP ring. Finally, a disulfide bond between TT residue C93 (β3-β4 loop) and MTP residue C79 (β7-β8 loop) links the connector to the tail tube (Fig. 3e).

### A disulfide-cross linked tail tube

The tail tube, composed of 58 stacked MTP hexamers, connects the TT ring at its proximal end to the distal tail protein (Dit) ring of the baseplate at its distal end (Fig. 1a,b). To overcome the flexibility of the long ФJabs tail, we reconstructed the three MTP rings immediately above the baseplate, which form a straight segment of the tube, combining particle re-extraction and localized refinement, yielding a 3.4 Å reconstruction (Fig. 4a, and Supplementary Fig. S2, S4, and Table S1). The full-length MTP consists of 352 amino acid residues (Fig. 1c and Supplementary Table S2). Although its central domain was well resolved, density for the peripheral domain was limited to a single α-helix and a few fragmented stretches (Fig. 4a).

**Fig. 4.**
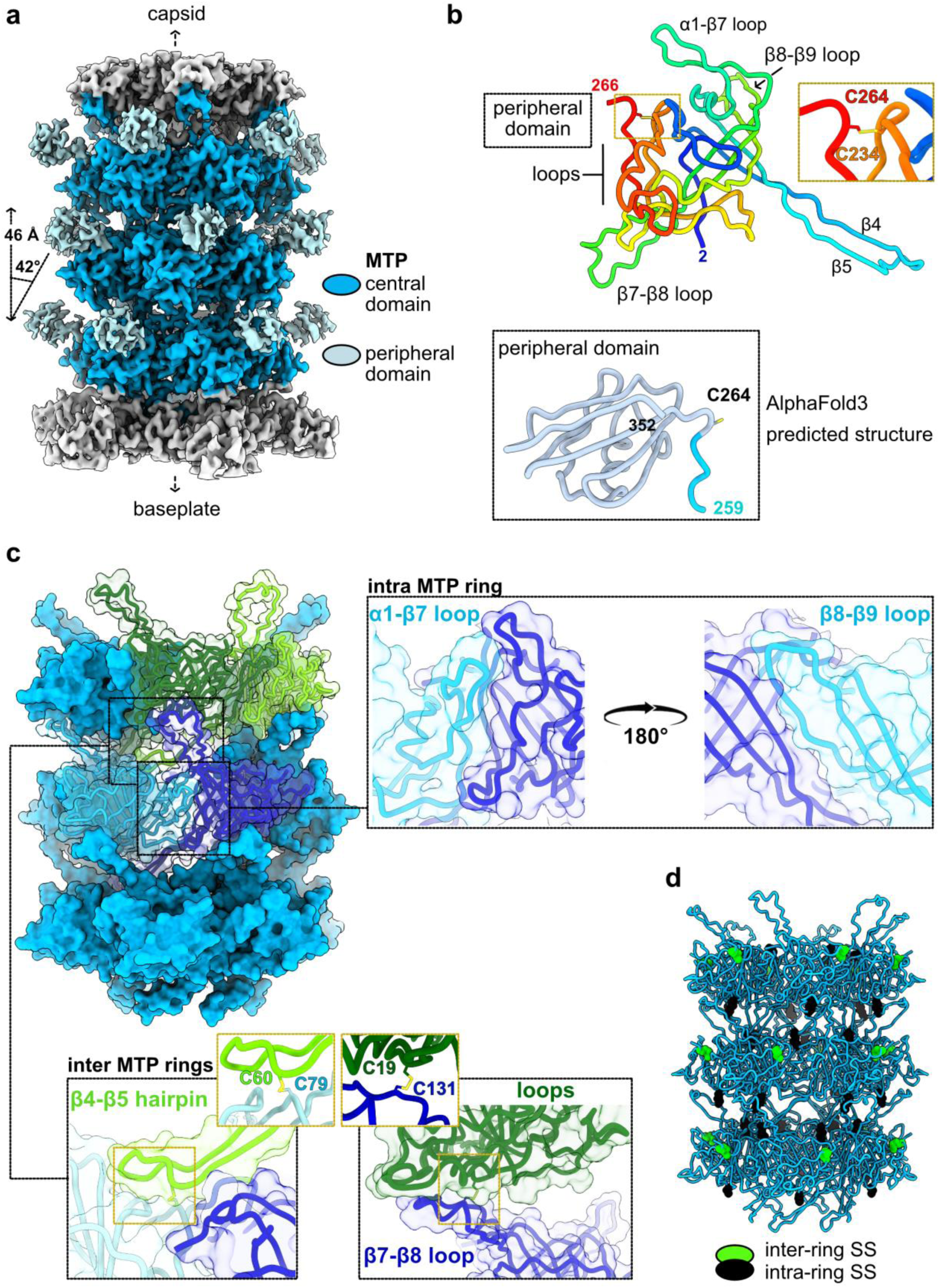
A disulfide-cross linked tail tube. **a.** 3D reconstruction of the tail tube segment bound to the baseplate. The three well-resolved MTP rings are colored. **b.** Rainbow-colored ribbon representation of the MTP central domain. Inset: ribbon representation of the AlphaFold3-predicted peripheral domain. **c.** Ribbon and surface representations of the three MTP rings. Insets show close-up views of intra- and inter-ring protein-protein interactions. Disulfide bonds are shown as sticks. **d.** Intra- and inter-ring disulfide bonds (SS) (green and black spheres, respectively) mapped onto the three MTP rings.

The MTP central domain (residues 2-266) comprises a β-sandwich formed by three- and five-stranded β-sheets, a single α-helix and six extended loops, including the prominent β4- β5 hairpin (Fig. 4b). (residues 265-352) consists of a second β-sandwich formed by two- and four-stranded β-sheets, together with a single α-helix (Fig. 4b, and Supplementary Fig. S7a). Whereas the central domain is structurally similar to the MTP of the phage lambda ^27^, the peripheral domain most closely resembles a domain of the minor capsid protein of the mycobacteriophage Patience ^28^ (Supplementary Table S3 and Fig. S7b,c). A disulfide bond between C234 and C264 at the junction of the two domains (Fig. 4b,d), resulting in a total of 348 bonds across the 58 MTP rings, both stabilizes the central domain and tethers the peripheral domain to the tail tube, restricting its mobility (Supplementary Table S4).

Successive MTP rings are separated by 46 Å and rotated by 42°. Within each hexamer, adjacent MTP subunits form an extensive interface primarily mediated by the α1-β7 and β8- β9 loops, the latter augmenting the five-stranded β-sheet of the adjacent subunit to form a continuous inter-subunit β-sheet (Fig. 4c). By contrast, contacts between successive rings are comparatively limited, likely underlying the intrinsic flexibility of the tail tube. These inter-ring contacts involve the β4-β5 hairpin, which reaches two MTP subunits in the underlying ring (toward the baseplate), and the β7-β8 loop, which contacts surface-exposed loops of a MTP subunit in the overlying ring (toward the capsid) (Fig. 4c). They are further stabilized by inter-ring disulfide bonds linking C60 (β4-β5 hairpin) to C79 of the underlying subunit and C131 (β7-β8 loop) to C19 of the overlying subunit (Fig. 4c,d), resulting in a total of 684 disulfide bonds across the 57 inter-ring interfaces. Together with the intra-ring disulfide bonds, the total number of disulfide bonds in the tail amounts to 1032 (table S4). Finally, the β4-β5 hairpin and β7-β8 loop adopt distinct conformations in the MTP rings adjacent to the Dit and TT, respectively, enabling specialized interactions with the baseplate and connector (Supplementary Fig. S7d). Accordingly, at the TT-MTP interface, the C93_TT_-C79_MTP_ disulfide bond replaces the C60_MTP_-C79_MTP_ bond (Fig. 3e, Fig. 4c), whereas at the MTP-Dit interface the C19_MTP_-C100_Dit_ bond replaces the C19_MTP_-C131_MTP_ bond (Fig. 4c, Fig. 5c, and Supplementary Table S4 and Fig. S8b).

**Fig. 5.**
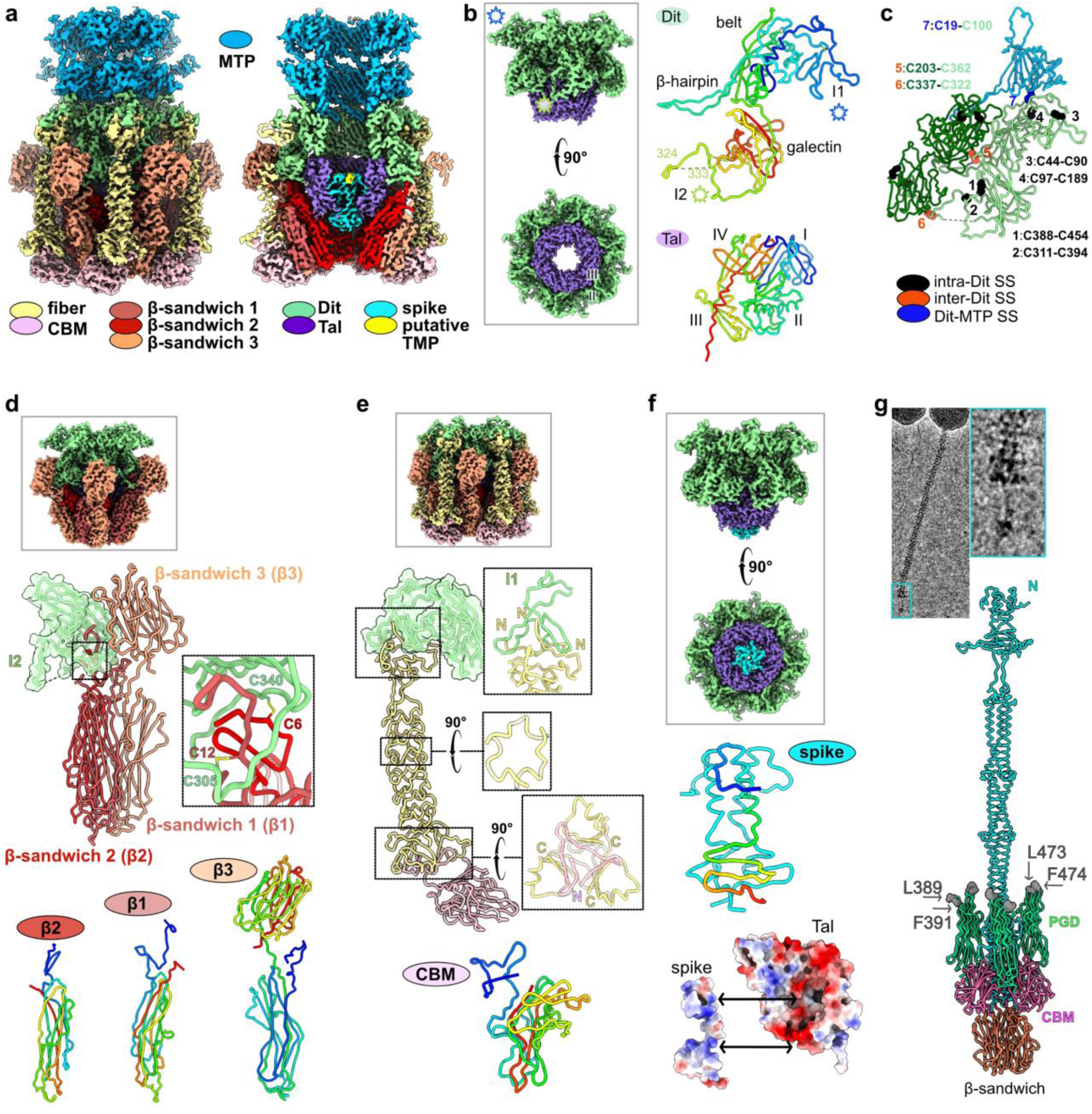
A complex baseplate with multiple potential receptor-binding modules. **a.** Cryo-EM reconstruction of the baseplate. The right panel shows a cut-away view highlighting interactions between the different components. **b.** (left) Reconstruction of the Dit-Tal central hub. (right) Ribbon representations of Dit and Tal. Stars indicate ФJabs Dit-specific structural features. **c.** Intra-Dit, inter-Dit, and Dit-MTP disulfide bonds (SS) (black, orange and blue spheres, respectively) mapped onto two adjacent Dit subunits and one MTP subunit. **d.** Reconstruction (top) and ribbon/surface representations (middle) of the Dit-bound β-sandwich heterotrimer. Inset: disulfide bonds linking β-sandwich 2 and β-sandwich 1 to Dit insertion I2. (bottom) Rainbow ribbon representations of the three proteins. **e.** Reconstruction (top) and ribbon/surface representations (middle) of the Dit-bound CBM-fiber complex. Insets: close-up view of the fiber trimer interactions with the Dit insertion I1 (top), cross-section of the fiber central rod (middle), and fiber trimer interactions with CBM. (bottom) CBM rainbow ribbon representation. **f.** (top) Reconstruction of the Dit, Tal and spike Nter region. (middle) Rainbow ribbon representation of the spike trimer (residues 2-61). (bottom) Electrostatic surfaces of Tal and spike highlighting charge complementarity. **g.** AlphaFold3 model of the full-length spike trimer. Micrographs highlight the arrow-like spike protruding from the baseplate. Surface-exposed hydrophobic residues in the polyglycine-rich domain (PGD) are shown as spheres.

### A baseplate with multiple potential receptor-binding modules

We reconstructed the baseplate together with the last MTP rings, applying C3 and C6 symmetry to reach overall resolutions of 3.7 and 3.3 Å, respectively (Fig. 1b, Fig. 5a, and Supplementary Table S1, Fig. S2 and Fig. S4). Its central core is a hexameric Dit ring attached to the tail tube on one face, and to the Tal trimer on the other. Three other components, which we named according to their fold and overall structure, decorate this hub: 1) six heterotrimers of β-sandwiches 1-3; 2) six trimers of the fiber protein each bound to a single carbohydrate-binding module (CBM); 3) a trimer of the spike protein (Figs. 1b,c, Fig. 5a, and Supplementary Tables S2 and S3).

#### The Dit-Tal central hub

We modeled most of the 478 residues composing the Dit protein (Fig. 1c and Supplementary Table S2) in the C6-symmetrized reconstruction, except for a short disordered loop (residues 325-333). Overall, the ФJabs Dit architecture resembles that of other siphophage Dit proteins and comprises the canonical N-terminal belt domain, an MTP-like fold characterized by an elongated β-hairpin (residues 149-178) that mediates hexamer assembly, and a peripheral galectin domain (Fig. 5b,c, and Supplementary Table S3 and Fig. S8a). The belt and galectin domains are stabilized by intra-molecular disulfide bonds (C44-C90, C97-C189, C311-C394 and C388-C454; Fig. 5b,c, and Supplementary Table S4 and Fig. S8b). The ФJabs Dit also contains two large insertions: one in the belt domain (residues 16-139; insertion I1) and the other in the galectin domain (residues 297-344; insertion I2). Insertion I1 forms a long peripheral loop, stabilized by the C44-C90 disulfide bond, that interacts with other components of the baseplate (Fig. 5b,c,d). Insertion I2 forms an extended loop, stabilized by the C311-C394 disulfide bond, that contacts the adjacent Dit subunit and forms an inter-subunit disulfide bond between C322 of one subunit and C337 of its neighbor. The Dit hexamer is further stabilized by inter-subunit disulfide bonds linking C203 (belt domain) of one subunit to C362 (galectin domain) of the adjacent subunit, resulting in a total of 36 bonds across the Dit ring (Fig. 5c, and Supplementary Table S4 and Fig. S8b) The Dit β-hairpin, galectin domain and insertion I2 all contact the underlying Tal trimer (Supplementary Fig. S8c). The ФJabs Tal (343 of 345 residues modeled, Table S2) comprises the four canonical structural domains found in several siphophages (e.g., lactic acid bacteria-infecting phages ^29^ ^30–32^), and its closest structural match is the Tal of the myophage Mu ^33^ (Fig. 5b, and Supplementary Table S3 and Fig. S8d). The N-terminal (domain I) and C-terminal (domain IV) β-sandwiches of the three Tal subunits assemble into a pseudo-hexameric structure that docks beneath the Dit hexamer (Supplementary Fig. S8e). At the distal end of the Tal trimer, domains II and III adopt an open conformation (Fig. 5b, and Supplementary Fig. S8e,i), creating a ∼20 Å-wide opening that is occupied by a trimer of the spike protein (described below).

#### Potential receptor-binding proteins anchored to the Dit-Tal hub

The periphery of the Dit-Tal core is covered by six heterotrimers of the β-sandwiches 1-3, themselves surrounded by six CBM-fiber complexes (Fig. 5a,c,d); a trimer of the spike protein is inserted within the Tal trimer.

#### The β-sandwich heterotrimer is reminiscent of a *Lactococcus lactis* phage receptor-binding protein

Six elongated heterotrimers of the β-sandwiches 1-3 decorate the periphery of the Dit-Tal core (Fig. 5a,d). Each is anchored to the baseplate through contacts with the Dit galectin domain and insertion I2. The C-terminal region of β-sandwich 3 (residues 208-426, 426 residues in total; table S2), which adopts a double β-sandwich fold, forms an intermolecular nine-stranded β-sheet with the Dit galectin domain, whereas the N-terminal regions of β-sandwich 2 (residues 2-20, 426 residues in total; table S2) and β-sandwich 1 (residues 3-30, 213 residues in total; table S2) insert into I2, forming the disulfide bonds C6_βsandwich2_-C340_Dit_ and C12 _βsandwich1_-C305_Dit_ (Fig. 5d, and Supplementary Table S4). The C-terminal region of β-sandwich 2 (residues 21-199) and β-sandwich 1 (residues 31-213), and the N-terminal domain of β-sandwich 3 (residues 3-207) share a similar β-sandwich fold composed of four- and three-stranded β-sheets that are bent at their center by β-bulges (Fig. 5c and Supplementary Fig. S8f). This fold resembles that of the receptor-binding domain of *L. lactis* phage 1358 ^30^ (Supplementary Table S3 and Fig. S8g), which recognizes saccharide receptors ^34^, suggesting that the β-sandwich heterotrimer may function as a receptor-binding module.

#### A CBM bound to a trimeric fiber protein

Each trimer of the fiber protein (residues 5-139 modeled, Table S2) forms a ∼110 Å-long rod anchored to the Dit insertion I1, which folds as a two-pronged clamp around the N-terminal end of the fiber trimer and contributes a three-stranded anti-parallel β-sheet with the N-terminal residues (6-9) of one fiber subunit (Fig. 5b,e). The predominantly loop-rich central regions of the three fiber subunits wrap around one another to form a helical rod with a star-shaped cross-section. At its distal end, the fiber trimer associates with the N-terminal region of the CBM (residues 1-34, Table S2) through the formation of intermolecular β-sheets. The CBM C-terminal region is similar to a glycosidase domain (Supplementary Table S3 and Fig. S8h), suggesting that the ФJabs CBM may interact with cell surface polysaccharides.

#### The trimeric spike contains three potential host-binding domains

We modeled three copies of the N-terminal region (residues 2-61) of the spike protein (788 residues; Supplementary Table S2). This trimeric assembly occupies the central cavity of the Tal trimer, with its distal end extending through and occluding the opening of the Tal channel (Fig. 5e). It comprises N-terminal 10-residue stretches, a three-helix bundle (residues 11-26), and a β-prism-like fold formed by the association of three three-stranded β-sheets, which contact Tal domains II, III and IV through complementary electrostatic surfaces (Fig. 5f). This binding mode resembles that described for the putative spike receptor-binding protein of the mycobacteriophage Badfish ^15^. Three density patches located in between the N-terminal stretches and helices may correspond to the TMP (Fig. 5a, and Supplementary Fig. S8I). Although an arrow-like extension protruding from the baseplate is clearly visible in micrographs (Fig. 1a, Fig. 5g), the corresponding density was not resolved in our 3D reconstructions centered on the Dit-Tal junction. AlphaFold3 predicts the full-length spike trimer to consist of the N-terminal Tal-anchoring module followed by an ∼250 Å-long helical extension that terminates in three distinct domains (Fig. 5g, and Supplementary Fig. S9a). The first is a polyglycine-rich domain similar to that of the cyanophage Pam3 ^35^ (Supplementary Table S3 and Fig. S9b). Similar to previously described mycobacteriophage polyglycine-rich domains, which display clusters of surface-exposed hydrophobic residues ^15^, the ФJabs polyglycine-rich domain exposes residues F389, L391, L473, and F474 in the loops at the tip of its polyglycine-helix bundle (Fig. 5g). These loops may contribute to host binding, as proposed for the polyglycine-rich domain of Salmonella phage S16 ^36^. The second and third domains are a CBM-like domain and an elongated β-sandwich domain, respectively, the latter forming the arrow-like tip of the spike trimer (Supplementary Fig. S9c,d). This trimeric β-sandwich architecture is reminiscent of receptor-binding domain trimers from phages infecting lactic acid bacteria (Supplementary Table S3 and Fig. S9d).

## DISCUSSION

The cryo-EM structure of ФJabs provides a high-resolution view of a phage originally selected on a *Mabs* clinical isolate ^4^. Beyond filling an important gap in our knowledge of mycobacteriophage architecture, this structure expands our understanding of phage structural diversity and provides structural insights for future investigations of the molecular determinants of host specificity, including recognition of the *Mabs* S and R variants. Among the growing number of high-resolution siphophage structures, ФJabs is distinguished by four remarkable architectural features: 1) a T=9 capsid assembled from two MCPs, a feature previously assigned to prolate capsids only ^37^; 2) a virion covalently linked from capsid to baseplate by an extensive network of ∼2,000 disulfide bonds; 3) a simplified three-component connector lacking a stopper protein; and 4) a complex baseplate comprising multiple candidate receptor-binding proteins, including uncommon heterotrimeric assemblies of β-sandwich domains.

### Building a T=9 icosahedral capsid with two MCPs

ФJabs assembles a T=9 icosahedral capsid, an increasingly documented architecture ^12,38,39^, using two distinct MCPs: MCPh exclusively forms the hexons, whereas MCPp is restricted to the pentons. In previously characterized T=9 phages ^12,38,39^, a single MCP forms both pentons and hexons, whereas T4 employs two MCPs but assembles a prolate rather than a T=9 capsid ^37^. This ФJabs dual-MCP strategy and structural specialization of MCPh and MCPp are consistent with the distinct geometric constraints imposed by the hexameric and pentameric environments: MCPp contains a bent spine helix and a reduced axial domain adapted to the higher curvature at the fivefold vertices, whereas MCPh possesses a C-terminal extension whose six copies assemble into the prominent hexon turret.

The most instructive comparison is with the *Mycobacterium smegmatis*-infecting phage Douge ^12^. Despite sharing an identical T=9 architecture composed of 11 pentons and 80 hexons, the two phages stabilize their capsids through fundamentally different strategies. Douge assembles its shell from a single MCP reinforced by 120 cement proteins bound at the icosahedral threefold axes, whereas ФJabs dispenses entirely with cement proteins. Instead, unlike most structurally characterized tailed phages, which rely on non-covalent interactions, cement proteins or isopeptide cross-links to stabilize their capsids ^21,40^, ФJabs employs an extensive network of intra- and inter-subunit disulfide bonds. Remarkably, extensive disulfide bonding is found not only within the capsid but also across the connector, tail, and baseplate, where it covalently links these components into a continuous assembly.

### Disulfide-cross-linking spanning the entire virion

An extensive network of disulfide bonds reinforces nearly every interface within the ФJabs virion. These covalent cross-links stabilize adjacent MCPs within and between capsomers, connect the tail terminator to the first tail tube ring, bridge successive MTP rings and the baseplate to the tail, and reinforce the baseplate by linking candidate receptor-binding proteins to the Dit hub. Such extensive covalent reinforcement by disulfide bonds is unusual but not unprecedented: disulfide bonds stabilize the capsids of the RNA phages Qβ ^41^ and PP7 by cross-linking coat protein dimers ^42^, covalently attach tail fibers to the baseplate in the cyanophage Pam3 ^35^, and spans the virion of the *Agrobacterium* myophage Milano ^43,44^ and gut siphophage PD491P1 ^45^.

In Milano, the disulfide-bond network increases the virion stability and has also been proposed to regulate sheath contraction through redox-dependent remodeling ^43,44^. In contrast, ФJabs possesses a non-contractile tail, eliminating the need to accommodate large conformational rearrangements during infection. Its disulfide bond network therefore may primarily function as a structural reinforcement, stabilizing the capsid and covalently integrating the connector, tail and baseplate into a continuous assembly. Disulfide bond formation requires an oxidizing environment, whereas virion assembly takes place in the reducing bacterial cytoplasm. This raises the intriguing possibility that the disulfide bonds network does not form during particle assembly within the cytoplasm but rather late in morphogenesis, during or after host cell lysis, and therefore may represent a maturation step that mechanically reinforces the infectious virion. Whether disulfide bond formation occurs spontaneously or is catalyzed by host- or phage-encoded oxidoreductases remains an open question.

### A simplified connector architecture

ФJabs adopts a simplified three-component connector architecture comprising only the portal, adaptor, and TT proteins, without an identifiable stopper protein. In most structurally characterized siphophages, the connector contains a stopper protein, as observed in the mycobacteriophages Douge and Bxb1 ^11,12,24,46,47^. In tailed phages, stopper proteins are proposed to seal the connector after DNA packaging, thereby contributing to genome retention and establishment of the head–tail interface ^24,48^. However, simplified connectors lacking a stopper protein have also been described in few other phages ^49,50^, indicating that the four-component connector architecture is not universally conserved among siphophages.

The absence of a stopper protein raises the question of how its canonical functions are achieved in ФJabs. The direct interaction between the adaptor and TT proteins suggests that these components may collectively contribute to functions normally associated with the stopper protein. Interestingly, in the mycobacteriophage Douge, the TMP extends through the stopper-containing connector and into the tail lumen ^12^, placing it along the DNA translocation channel. Although the connector architectures differ, this arrangement raises the possibility that the TMP may also participate in genome retention and/or release in ФJabs. Therefore, ФJabs illustrates the diversity of siphophage connector architectures, suggesting that functions traditionally associated with a dedicated stopper protein may be carried out by the remaining connector and tail components.

### A baseplate specialized for carbohydrate recognition?

The ФJabs baseplate is considerably more complex than those reported for many siphophages infecting Gram-positive or Gram-negative bacteria ^27,29,51,52^. The complexity of mycobacteriophage baseplates, in terms of both the number of candidate receptor-binding proteins and the diversity of their domain architectures and interactions with the Dit-Tal hub, was first revealed by our AlphaFold2-based analysis ^15^ and has since been confirmed by the cryo-EM structures of the *M. smegmatis* phages Bxb1 and Douge ^11,12^. A recent in-press publication describes the cryo-EM structure of the mycobacteriophage Mysterious baseplate ^9^. Although the authors report an overall topology and protein composition resembling those of ФJabs baseplate, detailed structural comparisons were not possible because the atomic coordinates and cryo-EM maps are not yet publicly available. Interestingly, the overall organization of the ФJabs baseplate also shares architectural features with those of *Streptococcus thermophilus* phages, including a long multidomain Tal extension reminiscent of the ФJabs Tal-spike assembly and peripheral receptor-binding protein trimers surrounding the central Dit-Tal hub ^53^, suggesting convergent evolution of multi-protein host-recognition machineries.

All three candidate receptor-binding proteins of ФJabs contain domains belonging to structural families associated with carbohydrate recognition. The β-sandwich heterotrimer resembles the saccharide-binding receptor-binding protein of the *L. lactis* phage 1358, the fiber trimer is associated to a CBM domain, and the distal spike carries a CBM domain. Together, these observations are consistent with the possibility that ФJabs recognizes carbohydrate-containing components of the mycobacterial cell envelope. Consistent with this hypothesis, ФJabs-resistant mutants characterized by a phage adsorption defect, harbor frameshift mutations in *MAB_2182c* encoding a putative glycosyltransferase ^4^, suggesting that glycosylation of an as-yet unidentified surface molecule is required for productive infection.

The preference of ФJabs for the GPL-rich S variant of *Mabs* further distinguishes it from previously characterized therapeutic *Mabs* phages, which preferentially infect the GPL-deficient R variant ^20^. These observations are consistent with the possibility that ФJabs exploits a receptor distinct from those recognized by R-specific phages, potentially involving GPL or another S-morphotype-associated surface glycoconjugate. Conversely, GPLs may mask the receptors targeted by R-specific phages. Moreover, whereas the glycolipid trehalose polyphleates act as a co-receptor for the therapeutic *Mabs* phages Muddy and BPs ^54^, they do not influence ФJabs infection, further supporting the existence of multiple receptor-recognition strategies among mycobacteriophages.

Lastly, the polyglycine-rich domain (PGD) of the spike protein presents surface-exposed hydrophobic residues, similar to previously identified mycobacteriophage polyglycine-rich domains ^15^, consistent with a potential role in interactions with lipid-containing components of the complex mycobacterial cell envelope ^55^. However, the lipid- and carbohydrate-binding activities inferred from the structure of the ФJabs baseplate remain to be experimentally validated. The cryo-EM structure presented here provides a foundation for the functional characterization of ФJabs host recognition and specificity, including the identification of its *bona fide* receptor-binding proteins and cognate cellular receptors.

## MATERIALS AND METHODS

### Phage production

ФJabs was amplified and purified as described in Liew JH et *al.* ^4^. The final phage sample was in SM buffer (50 mM Tris pH 7.5, 100 mM NaCl, 8 mM MgSO_4_).

### Phage genome extraction, sequencing, assembly and annotation

ФJabs genomic DNA (gDNA) was obtained using DNA extraction method as described in Liew JH et *al.* ^4^.

Genomic library for ФJabs was prepared using the Rapid Barcoding Kit (SQK-NBD114-24; ONT) and sequenced on a FLO-MIN114 flow cell using a MinION-Mk1B (ONT) for a running time of 72 h with default parameters.

The Command Line Application tool Dorado (v2.1.0; https://github.com/nanoporetech/dorado) was used to base-call and demultiplex the POD5 reads and output FASTQ files. For *de novo* assembly of ФJabs, Flye (v2.9.6) ^56,57^ was used. An online database, PhageScope (https://phagescope.deepomics.org) ^58^, was used for ФJabs annotation.

The resulting ФJabs assembly was inspected and manually curated to resolve structural protein coding regions required for cryo-EM modelling, namely the portal protein, tape measure protein (TMP) and tail-associated proteins. The consensus sequences from ФJabs genome assembly were then checked against the nucleotide database NCBI Basic Local Alignment Search Tool using BLAST to identify similar sequences and verify correctness.

### Cryo-EM grid preparation and data collection

2.5 µL of phage sample was applied to a 300-mesh copper Quantifoil R1.2/1.3 holey carbon grid previously glow-discharged for 60 sec at 30 mA using a GloQube system (Quorum Technologies). Following incubation for 10 sec at 20 °C and 100% relative humidity, the grid was blotted for 6.5 sec and vitrified in liquid ethane using a Vitrobot Mark IV (Thermo Fischer Scientific). Grids were imaged on a Glacios 2 transmission electron microscope (Thermo Fischer Scientific) operated at 200 kV, equipped with a Falcon 4i direct electron detector and Selectris-X energy filter, at the AFMB EM platform (Marseille, France). Movies were collected at 79,000 x yielding a pixel size of 0.9 Å, with a total dose of 40 e^-^/ Å ^2^ (40 frames/movie) and a defocus range of −0.8 to −2.4 µm.

### Cryo-EM image processing and single particle analysis

Single-particle image processing was performed in CryoSPARC v4.7.1 ^59^ to obtain 3D reconstructions of the ФJabs capsid, connector, part of the tail, and baseplate. Processing workflows and additional details are shown in figs. S1 and S2. Multi-frame movies were motion-corrected using Patch Motion Correction, and CTF parameters were estimated using Patch CTF. Capsid and adhesion-device particles were initially picked using blob picking, followed by template picking with 2D class averages generated from the blob-picked particles. Connector particles used for the final reconstruction were manually picked.

DNA-filled capsid particles were selected by 2D classification followed by *ab initio* reconstruction and heterogeneous refinement against two volumes with icosahedral (I) symmetry imposed. Selected particles were further refined by homogeneous refinement (I symmetry), yielding a 3.4 Å reconstruction used for atomic model building of the capsid asymmetric unit.

We first followed the procedure described by Hou et al. ^60^ to obtain a low-resolution 3D reconstruction of the connector. Capsid particles were symmetry expanded (I), and a cylindrical mask encompassing a fivefold vertex was generated in ChimeraX ^61^ for localized 3D classification (C1) to identify the unique portal vertex. The distance between the capsid center and the portal vertex was then used to re-extract particles centered on the capsid-portal junction. A low-resolution reconstruction generated from these particles served as the initial model for homogeneous refinement of manually picked connector particles with C5 symmetry imposed, allowing accurate alignment of the surrounding capsid density. Refined particles were subsequently C5 symmetry-expanded and subjected to localized 3D classification using a mask focused on the connector. Classes displaying well-resolved C12 and C6 structural features were combined and locally refined without symmetry (C1) using a mask encompassing the entire particle, yielding a 6 Å reconstruction. These tail-aligned particles were then C12- and C6-symmetry-expanded and locally refined using masks focused on the portal-adaptor and TT-MTP regions, producing reconstructions at 3.4 Å and 3.9 Å resolution, respectively. The two maps were combined to generate a composite reconstruction of the connector.

Adhesion-device particles were selected by 2D classification followed by *ab initio* reconstruction and heterogeneous refinement against two volumes with C6 symmetry imposed. Particles were then subjected to homogeneous refinement with C6 symmetry imposed, C6-symmetry expanded and 3D classified into five classes. Particles of the largest and best-defined 3D class (74.4% of particles, 4235 particles after duplicate removal) were used for two independent refinements: 1) C6 symmetry expansion followed by localized refinement using a mask encompassing the entire baseplate, yielding a 3.3 Å reconstruction; and 2) homogeneous refinement with C3 symmetry imposed, yielding a 3.7 Å reconstruction. The distance between the center of the baseplate and the second MTP ring was used to re-extract particles centered on a tail-tube segment. Following C6 symmetry expansion, localized refinement with masks encompassing first the entire particle and subsequently three MTP rings yielded a 3.4 Å reconstruction of the tail tube. Local resolution estimation was performed for all reconstructions in cryoSPARC ^59,62^ (figs. S3 and S4).

### Model building, refinement, validation and analysis

Structural models of the ФJabs virion proteins were generated using the AlphaFold3 server (https://alphafoldserver.com/) ^63^. Predictions were performed by taking into account the oligomeric state of each protein complex: MCPh hexamer, MCPp pentamer, dodecamers of the portal and adaptor proteins, hexamers of tail terminator and MTP, Dit hexamer, Tal trimer, β-sandwich heterotrimer, fiber trimers, and spike trimers. The CBM was predicted as a monomer. Low-confidence regions were removed from the predicted models, which were then rigid-body fitted into the cryo-EM maps using ChimeraX ^61^ and manually adjusted in Coot ^64^.

Model refinement, including atomic coordinates, occupancies, and B-factors, was performed in Phenix ^65^. Secondary structures were restrained, as well as Ramachandran and side-chain rotamers. Particular attention was paid to identifying and validating intra- and intermolecular disulfide bonds. Final models were validated in Phenix by assessing model geometry and map-to-model correlations. Structures were analysed using ChimeraX ^61^ and Coot ^64^. Structurally similar proteins were identified in the Protein Data Bank (PDB) and AlphaFold Protein Structure Database (AFDB) using Foldseek ^23^, and pairwise structural comparisons were performed using the DALI server ^66^.

## ACKNOWLEDGMENTS

We acknowledge Denis Ptchelkine, head of the EM facility at the AFMB laboratory (Marseille, France), where the cryo-EM data were collected (French Infrastructure for Integrated Structural Biology grant ANR-10-INSB-0005), as well as Sébastien Santini and Antoine Flautre for maintaining the computational infrastructure (France 2030 program grant ANR-21-ESRE-0052) supporting data processing at the Institut de Microbiologie de la Méditerranée, Marseille, France. Molecular graphics and analyses performed with UCSF ChimeraX, developed by the Resource for Biocomputing, Visualization, and Informatics at the University of California, San Francisco, with support from National Institutes of Health R01-GM129325 and the Office of Cyber Infrastructure and Computational Biology, National Institute of Allergy and Infectious Diseases.

## AUTHOR CONTRIBUTIONS

Conceptualization: A.G.; Methodology: C.C., A.G.; Validation: C.C., A.G.; Formal Analysis: C.C., B.S.Y.T., A.G.; Investigation: A.G.; Resources: J.H.L., P.B., L.K., A.G.; Data curation: J.H.L., B.S.Y.T., P.B.; Writing - Original Draft: C.C., A.G.; Writing - Review and Editing: J.H.L., P.B., L.K., C.C., A.G.; Visualization: C.C., A.G.; Supervision: A.G.; Project Administration: A.G.; Funding Acquisition: L.K., P.B., A.G.

## DATA, CODE, AND MATERIALS AVAILABILITY

The cryo-EM 3D reconstructions and the associated structural models have been deposited in the Electron Microscopy Data Bank and Protein Data Bank, respectively, under the following accession codes: EMD-59362 / PDB 33DD (capsid); EMD-59283 / PDB 32ZF (C12-symmetrized connector); EMD-59282 / PDB 32ZE (C6-symmetrized connector); EMD-59281 (connector with no symmetry imposed); EMD-59291 / PDB 32ZU (connector composite map); EMD-59397 / PDB 33EL (C3-symmetrized baseplate); EMD-59358 / PDB 33DA (C6-symmetrized baseplate); EMD-59284 / PDB 32ZG (tail tube). Raw Nanopore sequencing reads have been deposited in the NCBI Sequence Read Archive under BioProject accession number PRJNA1514955 and are publicly accessible at https://www.ncbi.nlm.nih.gov/bioproject/PRJNA1514955/. The updated and corrected genome sequence is available under GenBank accession number PQ001584.2; the previous version is PQ001584.1. The coordinates of the predicted ФJabs MTP monomer and spike trimer structures are available on Zenodo (10.5281/zenodo.24139454).

---

**Fig. S1.**
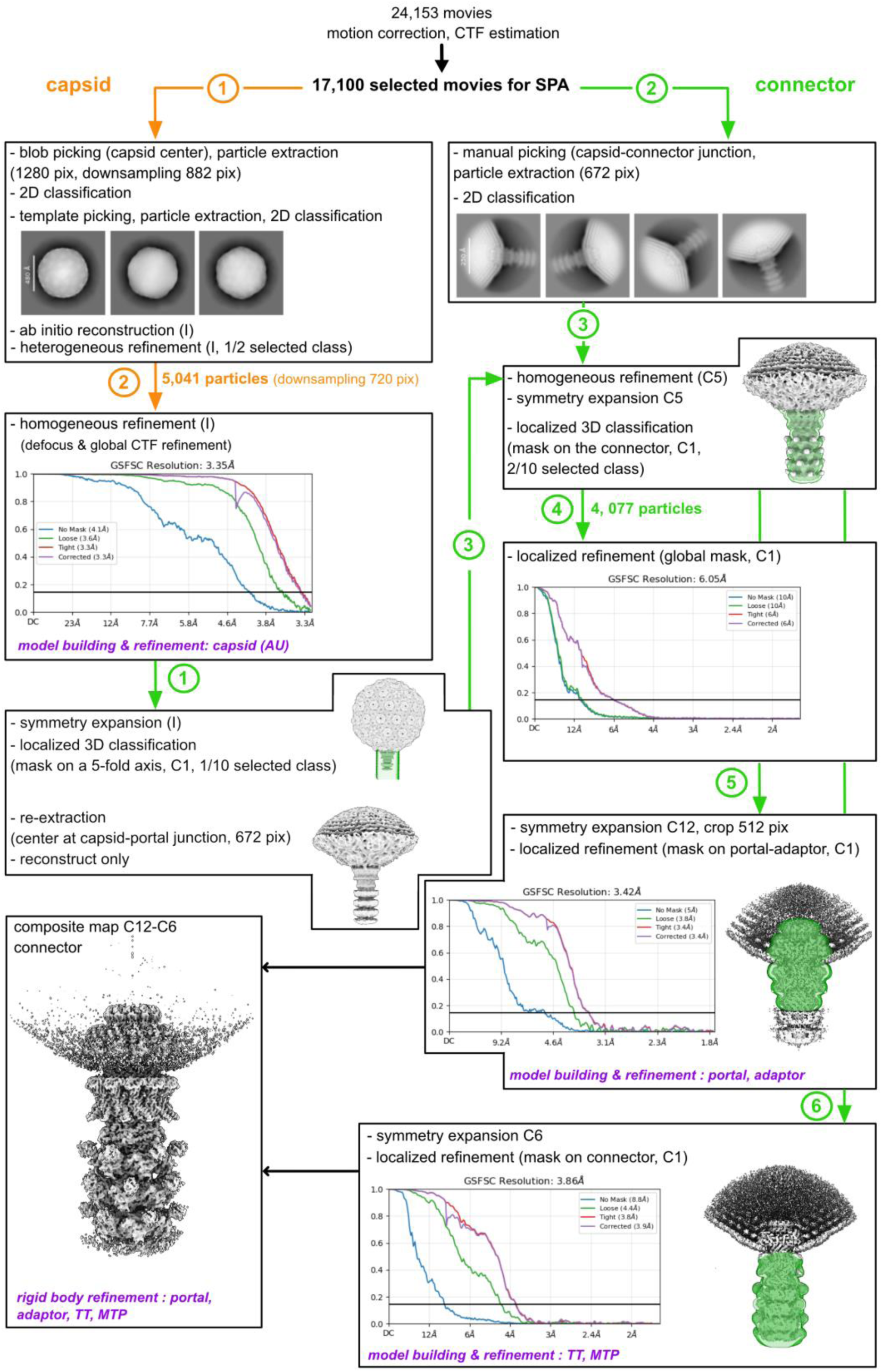
CryoEM structure determination workflow for the capsid and connector. Processing steps of the capsid and connector are shown in orange and green, respectively. Masks used for 3D classification and localized refinement, and gold-standard FSC curves (cut-off 0.143) are shown. Model building and refinement is indicated in purple.

**Fig. S2.**
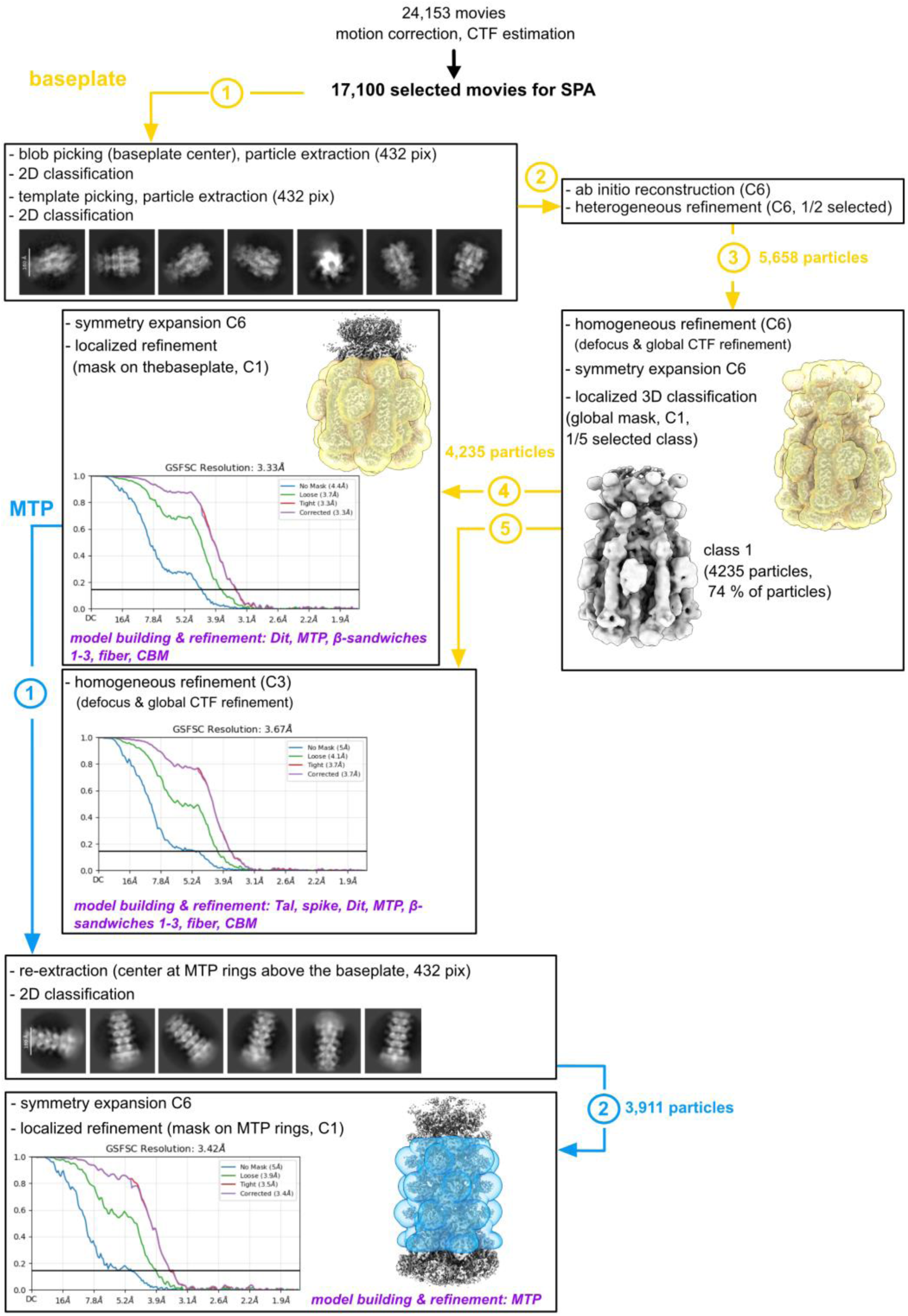
CryoEM structure determination workflow for the baseplate and MTP. Processing steps for the baseplate and MTP are shown in yellow and blue, respectively. Masks used for 3D classification and localized refinement, and gold-standard FSC curves (cut-off 0.143) are shown. Model building and refinement is indicated in purple.

**Fig. S3.**
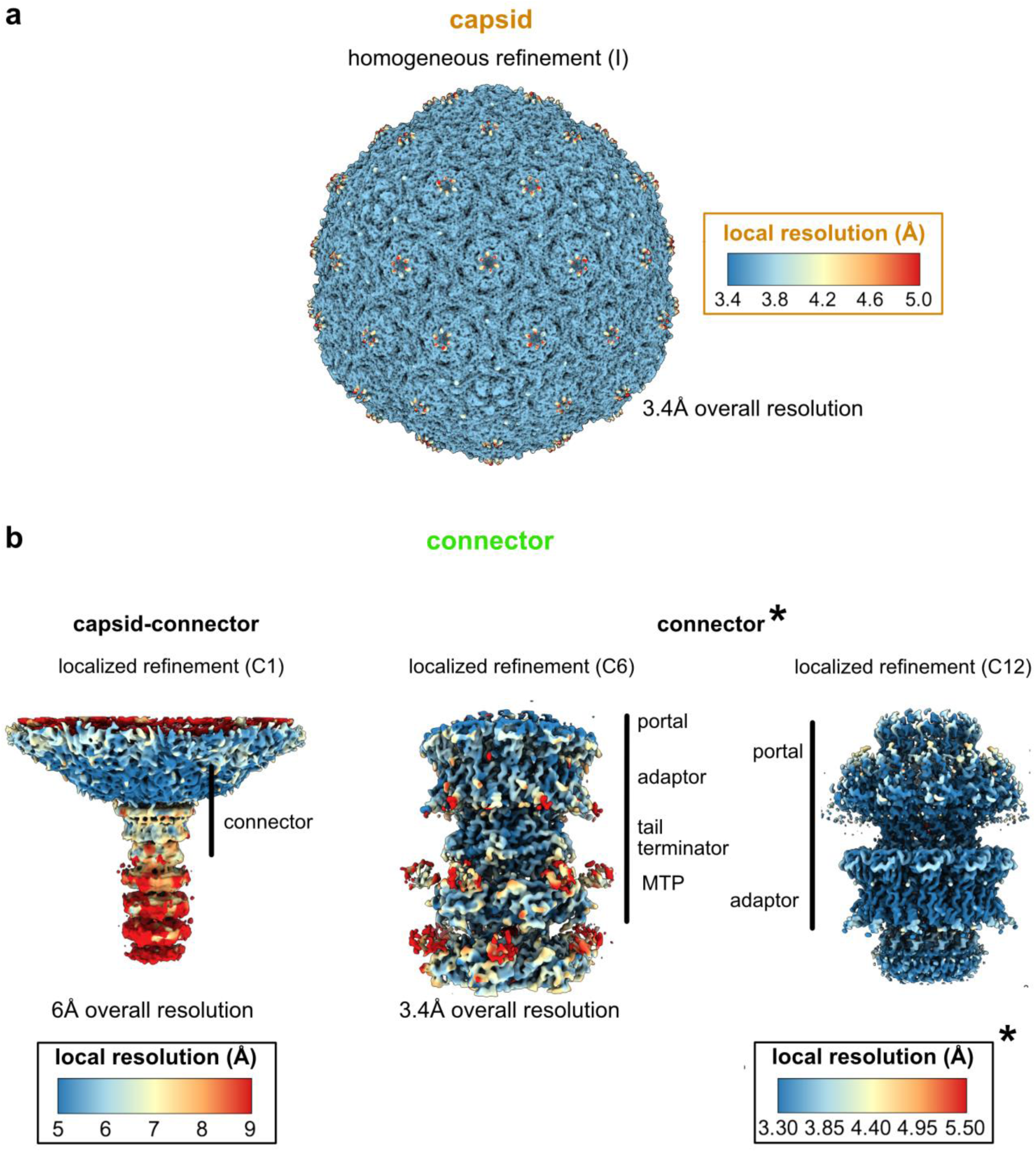
Local resolution distribution of the capsid and connector cryoEM maps. Surface representations colored according to local resolution are shown for the (**a**) capsid and (**b**) connector. The overall resolution and refinement strategy for each map are indicated in the corresponding view; the complete information is provided in Supplementary Fig. S1 and Table S1. The minimum value and range of each color scale were selected based on the overall resolution of the corresponding map to highlight meaningful local variations in resolution.

**Fig. S4.**
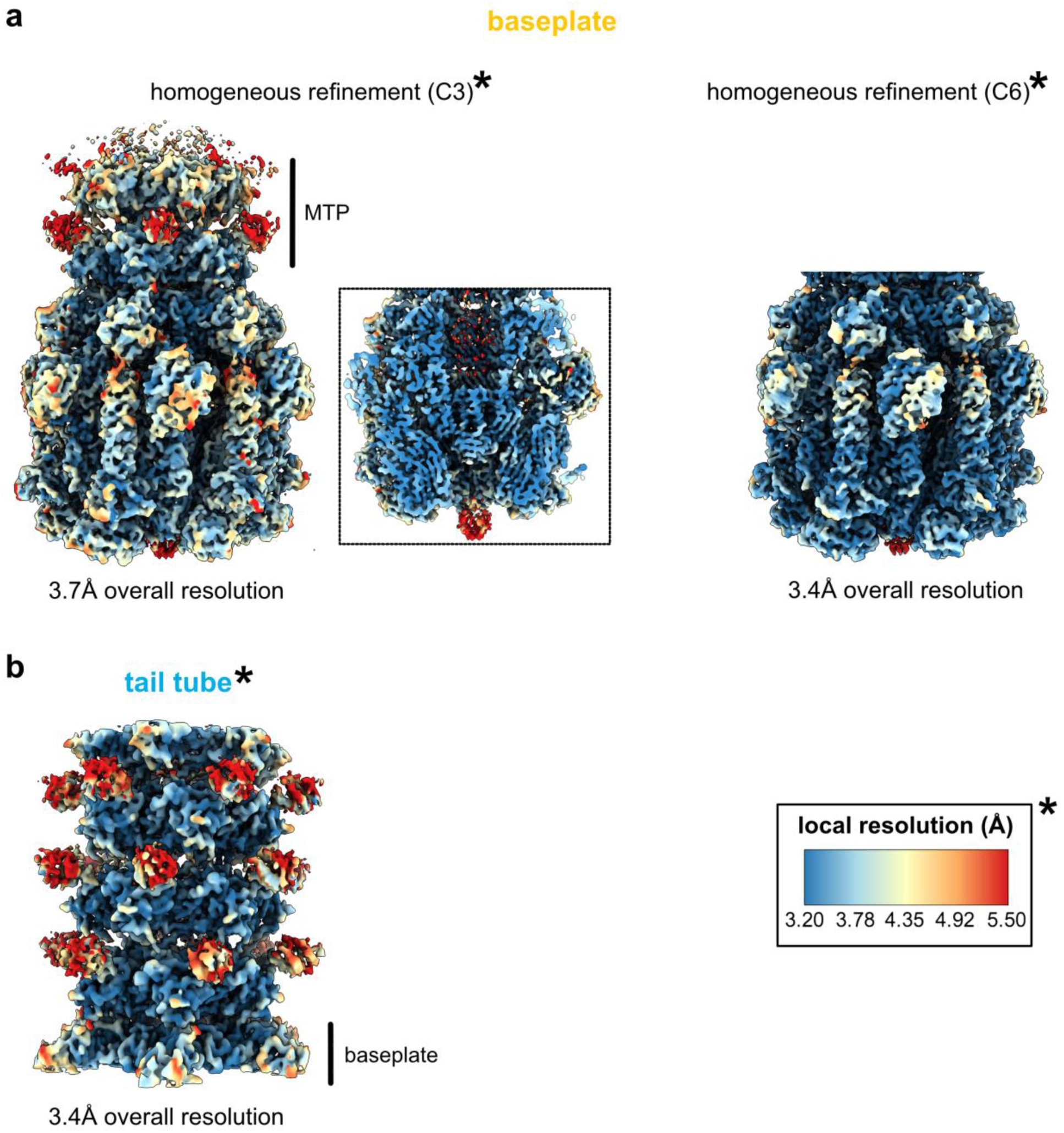
Local resolution distribution of the baseplate and tail tube cryoEM maps. Surface representations colored according to local resolution are shown for the (**a**) baseplate and (**b**) tail tube. A cut-away view of the C3-symmetrized baseplate reconstruction, indicated by the dotted box, reveals the internal density. The overall resolution and refinement strategy for each map are indicated in the corresponding view; the complete information is provided in Supplementary Fig. S2 and Table S1. The minimum value and range of each color scale were selected based on the overall resolution of the corresponding map to highlight meaningful local variations in resolution.

**Fig. S5.**
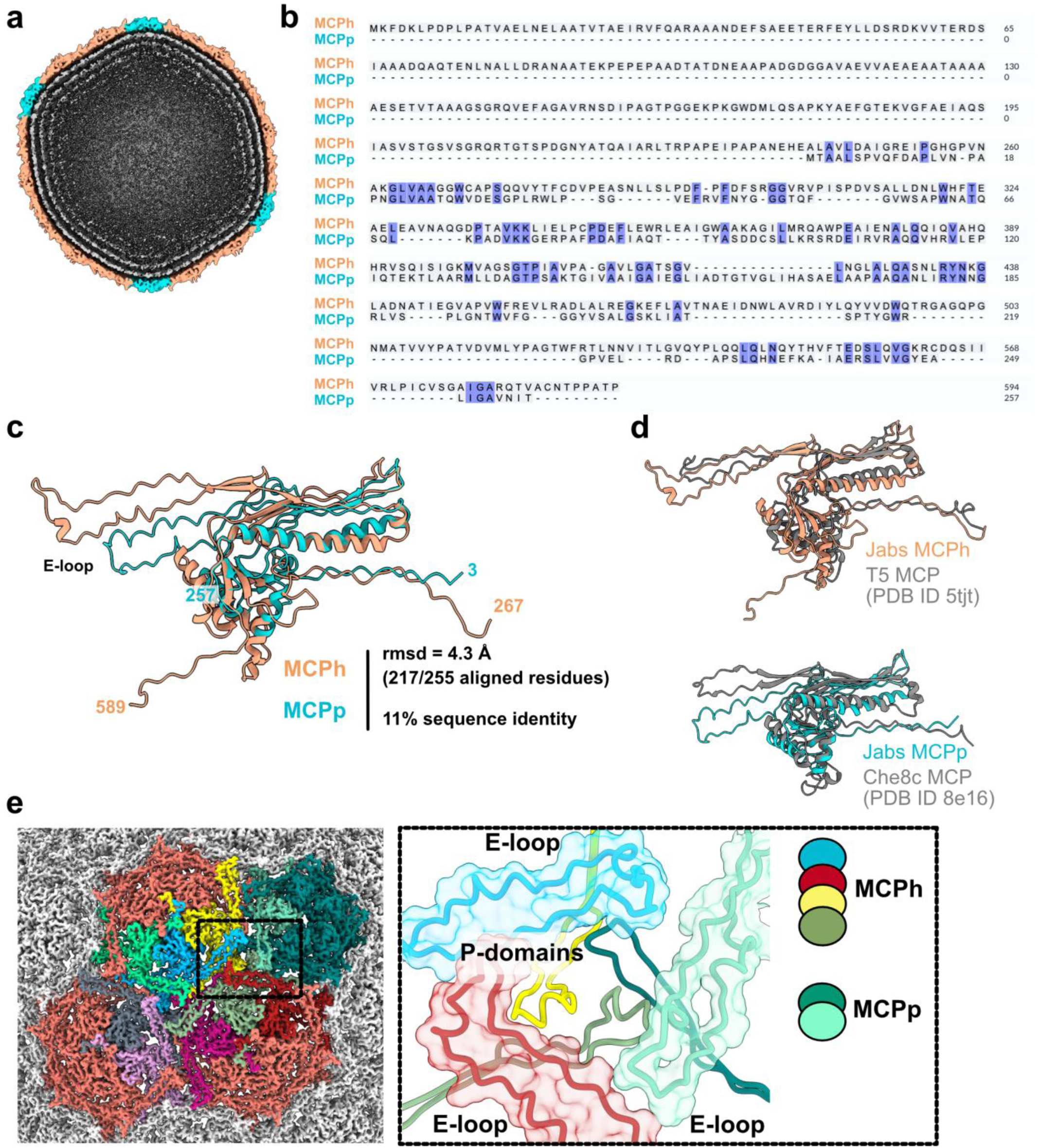
The ФJabs capsid is formed by two MCPs. **a.** Entire (top) and cut-away (bottom) views of the capsid reconstruction showing the asymmetric unit and concentric rings of the phage dsDNA genome, respectively. **b.** MCPh and MCPp sequence alignment (https://www.ebi.ac.uk/jdispatcher/msa/clustalo). **c.** MCPh and MCPp structure superposition. **d.** Structural superposition of the ФJabs MCPh (top) and MCPp (bottom) with the corresponding hits retrieved from Foldseek. **e.** (left) Reconstruction with MCPh and MCPp subunits highlighted to illustrate inter-capsomer interactions. (right) Close-up of the junction of two hexons and one penton showing E-loops (ribbons and surfaces) and central P-domains (ribbons only). The MCPp E-loop conformation disrupts the local 3-fold symmetry.

**Fig. S6.**
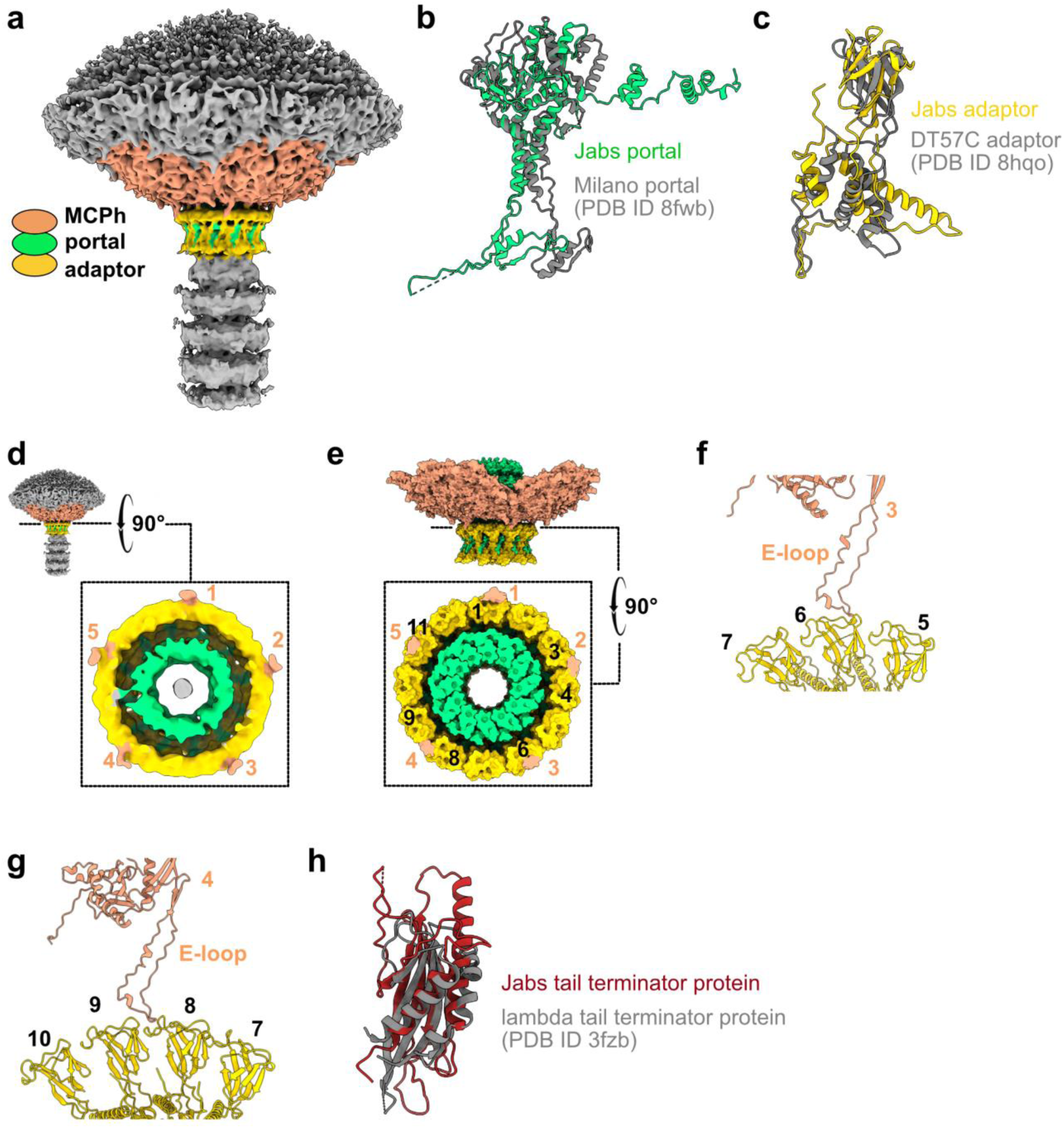
The ФJabs connector is formed by the portal, adaptor and tail terminator proteins. **a.** Asymmetric capsid-connector 3D reconstruction. The MCPh subunits surrounding the connector, as well as the portal and adaptor subunits, are colored. **b-c.** Structural superposition of the ФJabs portal (b) and adaptor (c) with the corresponding hits retrieved from Foldseek. **d.** Close-up view of the adaptor-capsid contacts in the asymmetric capsid-connector 3D reconstruction. **e.** Close-up view of the adaptor-capsid contacts in the MCPh-portal-adaptor structural model rigid-body fitted into the reconstruction. **f-g.** E-loops of MCPh subunits interacting with one (f) or two (g) adaptor subunits. **h.** Structural superposition of the ФJabs TT with the corresponding hit retrieved from Foldseek.

**Fig. S7.**
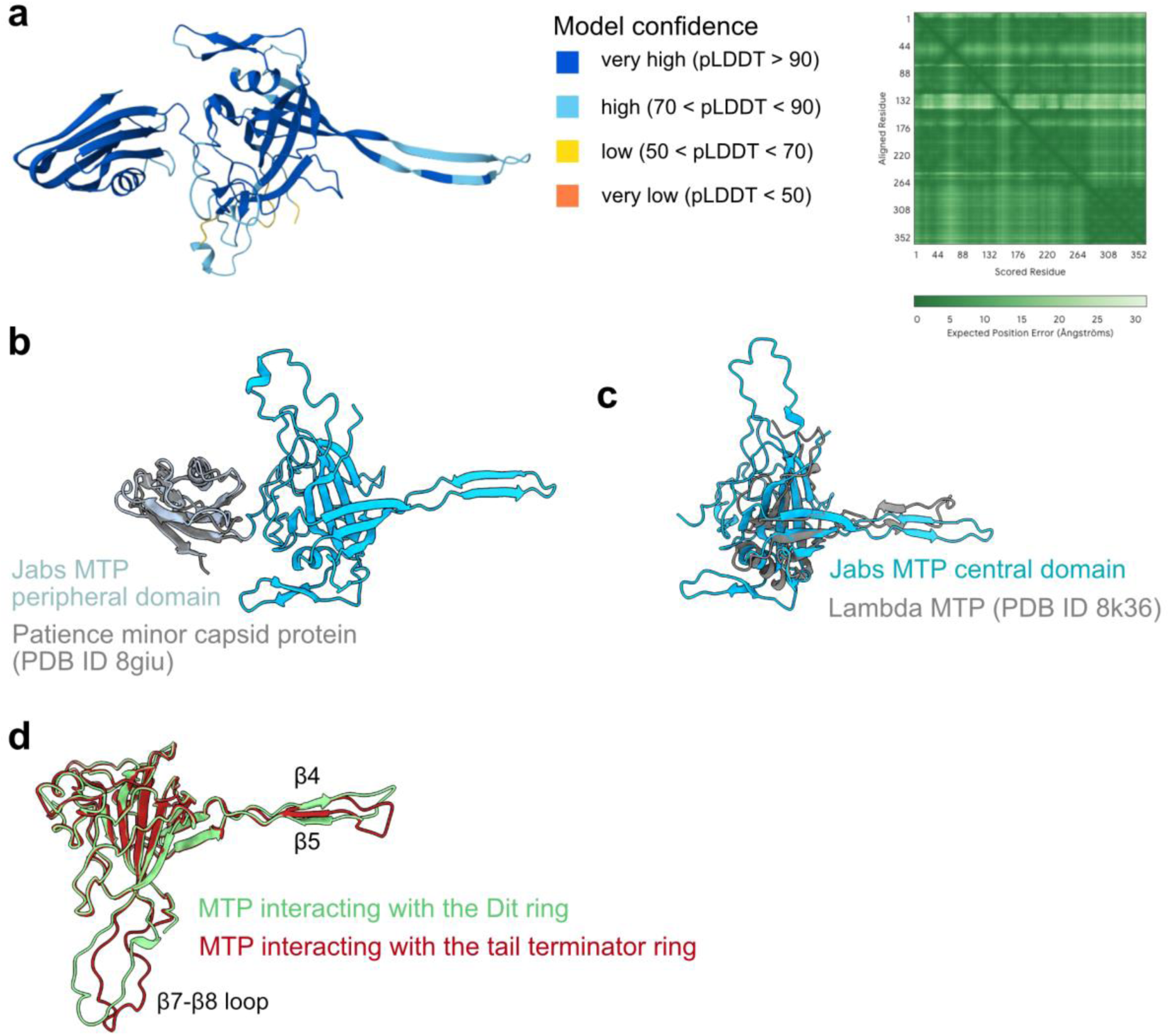
The ФJabs MTP contains a central domain and a peripheral domain. **a.** ФJabs MTP predicted structure by AlphaFold3. The structure is shown as ribbons colored according to the pLDDT values. The PAE plot is shown (right). **b-c.** Structural superposition of the ФJabs MTP with the corresponding hits retrieved from Foldseek for the peripheral (B) and central (C) domains. **d.** Structural superposition of the ФJabs MTP subunits interacting with the tail terminator ring (dark red) and the Dit ring (green).

**Fig. S8.**
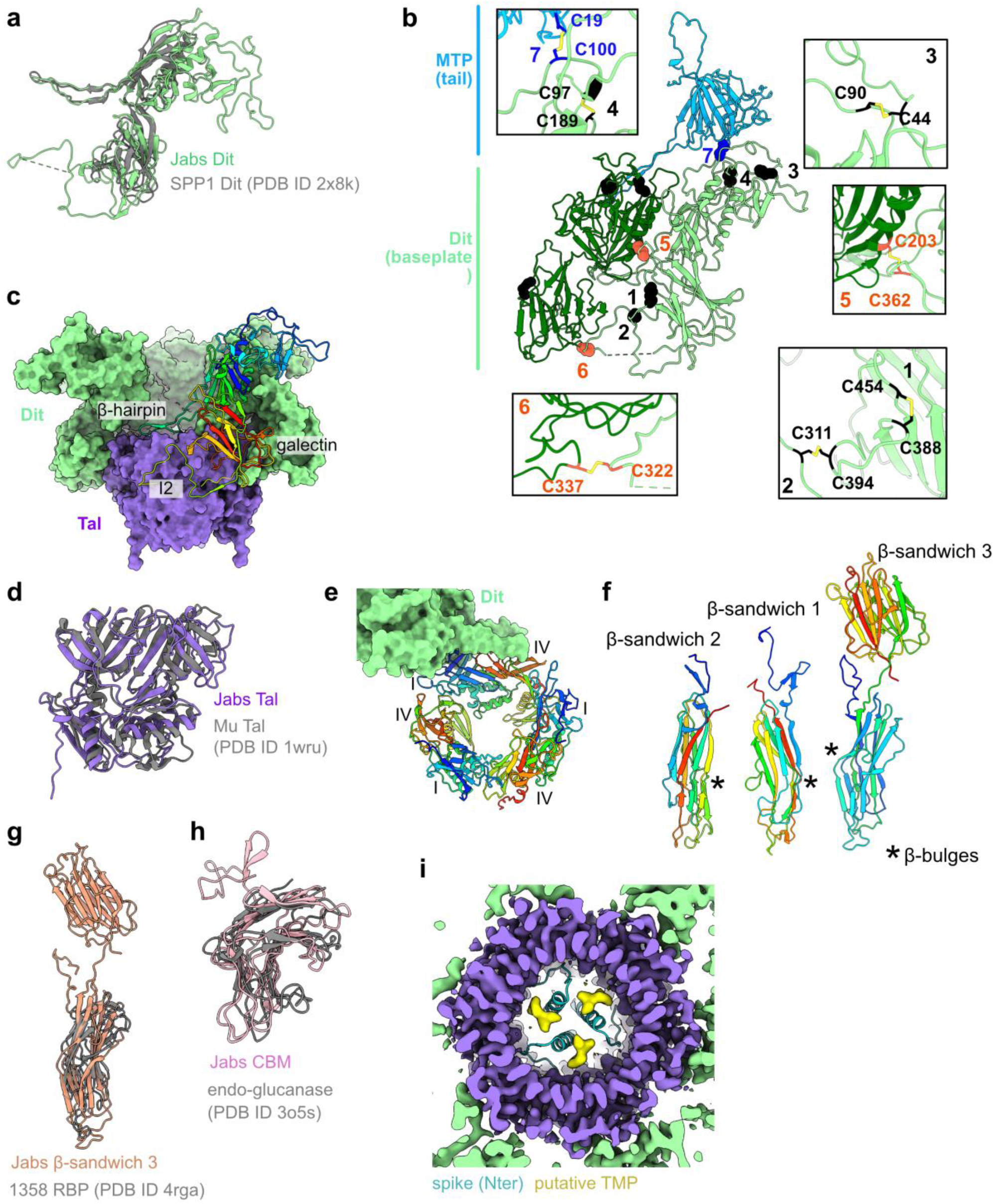
The ФJabs baseplate is a multiprotein assembly. **a.** Structural superposition of the ФJabs Dit with the corresponding hit retrieved from Foldseek. **b.** Intra-Dit, inter-Dit and Dit-MTP disulfide bonds (black, orange and blue spheres, respectively) mapped onto two adjacent Dit subunits and one MTP subunit. Insets: close-up views on ribbon representations of the proteins highlighting the disulfide bonds shown as sticks. **c.** Ribbon and surface representations of the Dit-Tal assembly highlighting the Dit regions in interaction with the Tal trimer. A Dit subunit is not shown for clarity. **d.** Structural superposition of the ФJabs Tal with the corresponding hits retrieved from Foldseek. **e.** Ribbon and surface representations of the Tal trimer (rainbow) and part of a Dit subunit highlighting the Tal domains in interaction with one Dit subunit. **f.** Rainbow ribbon representations of β-sandwiches 1-3. **g-h.** Structural superposition of the ФJabs β-sandwich 1 (g) and CBM (h) with the corresponding hits retrieved from Foldseek. **i.** Close-up view of the baseplate reconstruction showing three densities (yellow) within the Tal cavity, which may correspond to the TMP.

**Fig. S9.**
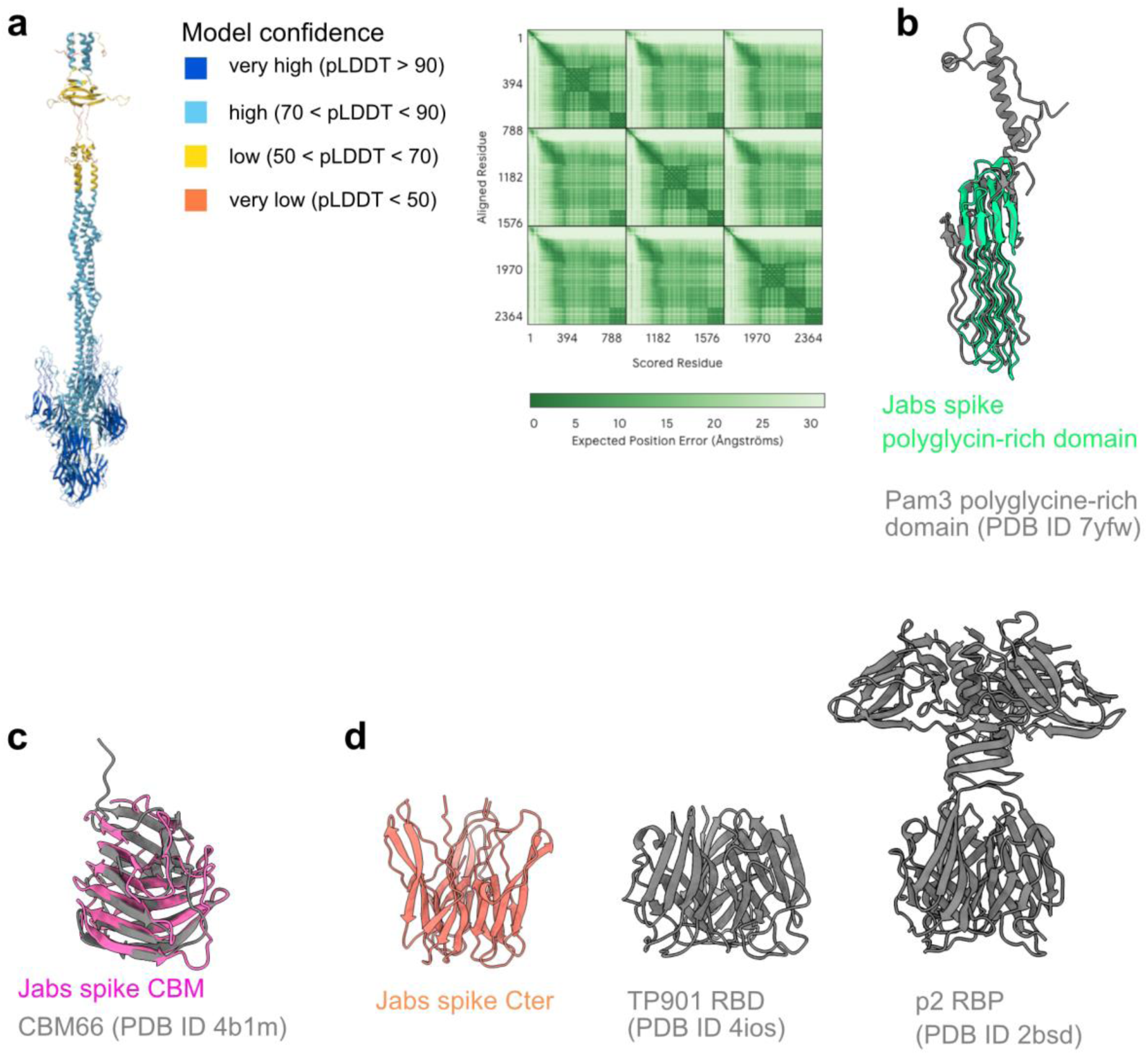
The ФJabs spike protein predicted structure. **a.** ФJabs spike trimeric structure predicted by AlphaFold3. The structure is shown as ribbons colored according to the pLDDT values. The PAE plot is shown (right). **b-c.** Structural superposition of the ФJabs spike polyglycine-rich domain (b) and CBM domain (c) with the corresponding hits retrieved from Foldseek. **d.** Structures of the ФJabs spike C-terminal domains (left), the *Lactococcus* phage TP901 receptor-binding domains, and the *Lactococcus lactis* phage p2 receptor-binding proteins (right) highlight their structural similarity.

**Table S1.** CryoEM 3D reconstruction and model refinement statistics (TT: tail terminator, MTP: major tail protein, CBM:carbohydrate-binding module).

| **3D reconstructions** | **Capsid** | **Connector** |  |  |  | **Baseplate** |  | **Tail tube** |
|---|---|---|---|---|---|---|---|---|
| Symmetry imposed | I | C12 | C6 | C1 | Composite map<br>C12-C6 | C3 | C6 | C6 |
| No. final particles | 5041 | 4077 |  |  | - | 4235 |  | 3911 |
| Resolution<br>(Å, FSC threshold 0.143) | 3.4 | 3.4 | 3.9 | 6.0 | - | 3.7 | 3.3 | 3.4 |
| Sharpening B factor (Å<sup>2</sup>) | 79.2 | 60.7 | 39.7 | 27.7 | - | 29.2 | 42.4 | 36.9 |
| EMDB code | 59362 | 59283 | 59282 | 59281 | 59291 | 59397 | 59358 | 59284 |
| **Structural models** | **capsid (AU)** | **portal, adaptor** | **TT, MTP** | **-** | **portal, adaptor, TT, MTP** | **Tal, spike (Nter), Dit, MTP, β-sandwiches 1-3, CBM, fiber** | **Dit, MTP, β-sandwiches 1-3, CBM, fiber** | **MTP** |
| No. chains | 10 | 24 | 12 | - | 36 | 60 | 54 | 18 |
| No. protein residues | 2839 | 7572 | 2634 | - | 10206 | 14412 | 13206 | 4770 |
| No. atoms | 21690 | 56345 | 19167 | - | 75512 | 108550 | 99398 | 35816 |
| Ligands | SO<sub>4</sub><sup>2-</sup> (1) | - | - | - | - | - | - | - |
| B factors (Å<sup>2</sup>)<br>protein<br>ligand | 72.2<br>97.5 | 84.6<br>- | 131.5<br>- | -<br>- | 108.2<br>- | 100.5<br>- | 86<br>- | 91.2<br>- |
| r.m.s.d.<br>bond lengths(Å)<br>bond angles (°) | 0.003<br>0.64 | 0.002<br>0.53 | 0.003<br>0.64 | - | 0.003<br>0.56 | 0.003<br>0.63 | 0.003<br>0.62 | 0.004<br>0.74 |
| Rotamers outliers (%) | 0 | 0 | 0 | - | 0 | 0 | 0 | 0 |
| Ramachandran plot (%)<br>favored<br>allowed<br>outliers | 94.2<br>5.8<br>0 | 95.1<br>4.9<br>0 | 94.3<br>5.7<br>0 | -<br>-<br>- | 94.9<br>5.1<br>0 | 94<br>6<br>0 | 94.1<br>5.9<br>0 | 96.5<br>3.5<br>0 |
| Molprobity score | 1.5 | 1.79 | 2 | - | 1.9 | 1.62 | 1.58 | 1.6 |
| Clashscore | 3.5 | 8.2 | 12.5 | - | 10.5 | 4.4 | 4 | 6.8 |
| Model-to-map fit<br>CC (ligand) | 0.88<br>(0.84) | 0.83 | 0.83 | - | 0.83 | 0.83 | 0.86 | 0.86 |
| PDB code | 33DD | 32ZF | 32ZE | - | 32ZU | 33EL | 33DA | 32ZG |

**Table S2.** Genbank accession numbers and amino acid lengths of proteins identified in the ФJabs virion structure.

| **Protein** | **Genbank accession number** | **n. amino acids** |
|---|---|---|
| Portal protein* | YGA28996* | 617 |
| major capsid protein (hexon; MCPh) | XES72125 | 594 |
| major capsid protein (penton; MCPp) | XES72126 | 257 |
| major tail protein (MTP) | XES72130 | 352 |
| adaptor protein | XES72131 | 247 |
| tail terminator | XES72132 | 180 |
| tape measure protein (TMP)* | YGA28997* | 1863 |
| distal tail protein (dit) | XES72139 | 478 |
| Tail-associated lysin (Tal) | XES72140 | 345 |
| spike protein* | YGA28998* | 788 |
| β-sandwich 1 | XES72115 | 213 |
| β-sandwich 2† | YGA28999† | 199 |
| β-sandwich 3 | XES72113 | 426 |
| fiber protein | XES72110 | 140 |
| carbohydrate-binding module (CBM) | XES72109 | 232 |

\*: genes for which previously split-gene annotations have been revised in the updated GenBank deposition (PQ001584.2).
† : gene for which the sequence has been revised in the updated GenBank deposition (PQ001584.2).

**Table S3:** Similarity between the structures of the ФJabs virion components and PDB deposited structures.

|  | **Foldseek** |  |  | **DALI pairwise alignment** |  |  |
|---|---|---|---|---|---|---|
| **Protein** | **Hit** | **E-value** | **PDB Id** | **Seq id. %** | **rmsd Å** | **n. aligned residues** |
| **MCPh** | *Escherichia coli* phage T5 MCP | 3 e-6 | 5tjt | 10.4 | 6.8 | 254/323 |
|  | mycobacteriophage Che8 MCP | 2.45 e-5 | 8e16 | 7.6 | 3.6 | 242/323 |
| **MCPp** | mycobacteriophage Che8 MCP | 5.7 e-7 | 8e16 | 16.3 | 4.2 | 212/255 |
| **Portal** | *Agrobacterium* phage Milano portal | 3.7 e-9 | 8fwb | 10.8 | 5.9 | 271/426 |
| **Adaptor** | *E. coli* phage DT57C adaptor | 3.0 e-5 | 8hqo | 17 | 4.9 | 99/203 |
| **TT** | *E. coli* phage lambda TT | 7.1 e-5 | 3fzb | 11.2 | 2.9 | 120/174 |
| **MTP** |  |  |  |  |  |  |
| 1-264 | *E. coli* phage lambda MTP | 5.8 e-3 | 8k36 | 9.5 | 3.7 | 131/263 |
| 265-352 | mycobacteriophage Patience<br>minor capsid protein | 4.4 e-3 | 8giu | 24.1 | 1.5 | 82/88 |
| **Dit** | *Bacillus subtilis* phage SPP1 Dit | 1.6 e-8 | 2x8k | 11.8 | 4.0 | 231/467 |
| **Tal** | phage Mu Tal | 1.3 e-14 | 1wru | 16.9 | 3.2 | 280/342 |
| **CBM** | Endo-glucanase | 7.1 e-5 | 3o5s | 11 | 3.1 | 44/60 |
| **fiber** | Mycobacterium | 1.4 e-10 | *AFDB*<sup>£</sup> | 46.5 | 1.2 | 164/231 |
| **β-sandwich 3** | *Lactococcus lactis* phage 1358 RBP | 2.1 e-3 | 4rga | 9.2 | 4.4 | 130/202 |
| **spike** |  |  |  |  |  |  |
| Nter (2-61) | Mycobacterium | 1.7 e-4 | *AFDB*<sup>%</sup> | 43.3 | 2.8 | 47/60 |
| Polyglycine-rich domain | Cyanophage Pam3 tail fiber | 4.8 e-6 | 7yfw | 19.9 | 1.3 | 45/188 |
| CBM-like | CBM66 | 7.5 e-4 | 4b1m | 10.5 | 0.9 | 55/155 |

<sup>£</sup> AF-A0A5S9MN12-F1-model\_v6
<sup>%</sup> AF-A0A1X0B724-F1-model\_v6.

**Table S4:**
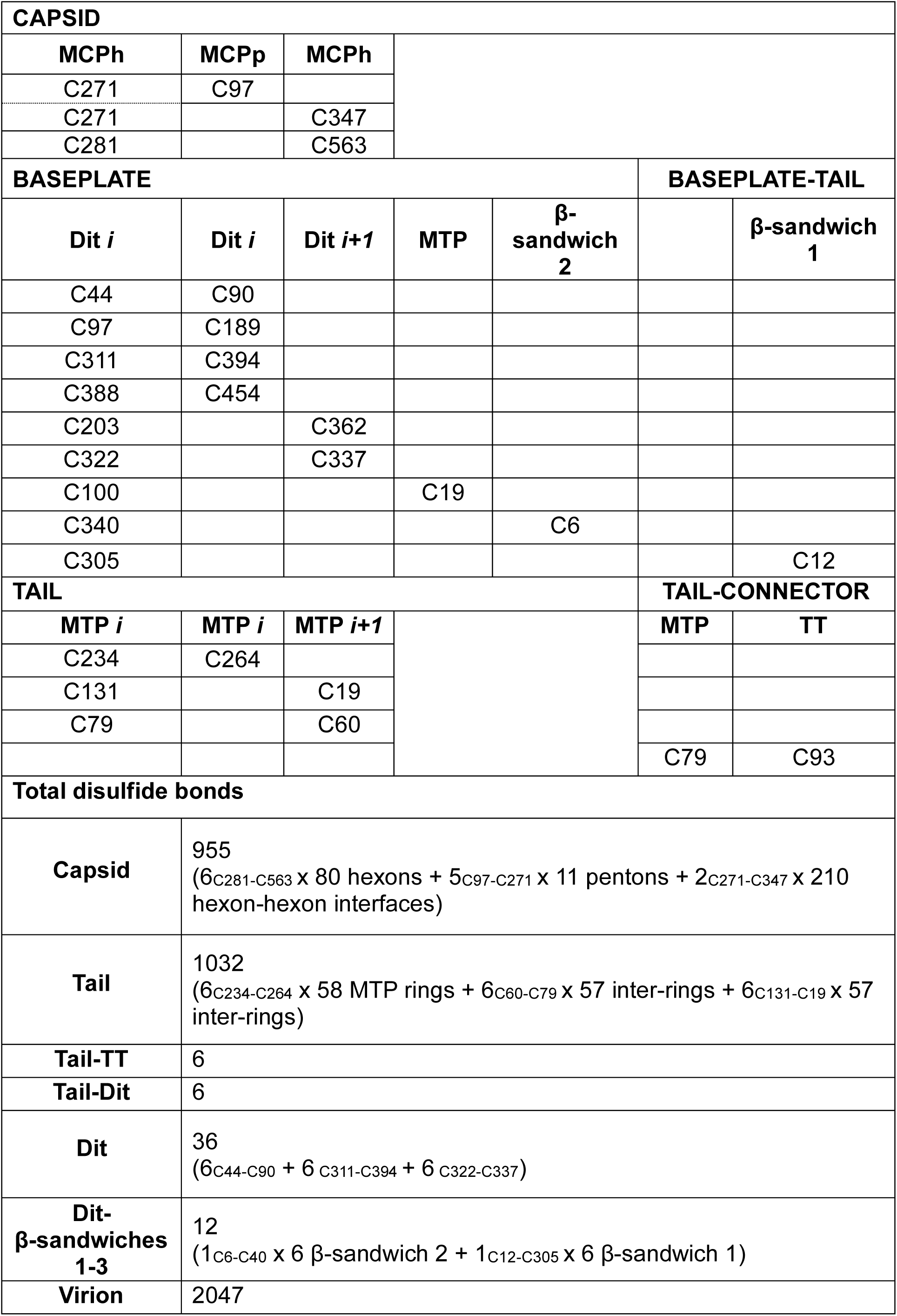
List of cysteine residues involved in disulfide bonds.

| **CAPSID** |  |  |  |  |  |  |
|---|---|---|---|---|---|---|
| **MCPh** | **MCPp** | **MCPh** |  |  |  |  |
| C271 | C97 |  |  |  |  |  |
| C271 |  | C347 |  |  |  |  |
| C281 |  | C563 |  |  |  |  |
| **BASEPLATE** |  |  |  |  | **BASEPLATE-TAIL** |  |
| **Dit *i*** | **Dit *i*** | **Dit *i+1*** | **MTP** | **β-sandwich 2** |  | **β-sandwich 1** |
| C44 | C90 |  |  |  |  |  |
| C97 | C189 |  |  |  |  |  |
| C311 | C394 |  |  |  |  |  |
| C388 | C454 |  |  |  |  |  |
| C203 |  | C362 |  |  |  |  |
| C322 |  | C337 |  |  |  |  |
| C100 |  |  | C19 |  |  |  |
| C340 |  |  |  | C6 |  |  |
| C305 |  |  |  |  |  | C12 |
| **TAIL** |  |  |  |  | **TAIL-CONNECTOR** |  |
| **MTP *i*** | **MTP *i*** | **MTP *i+1*** |  |  | **MTP** | **TT** |
| C234 | C264 |  |  |  |  |  |
| C131 |  | C19 |  |  |  |  |
| C79 |  | C60 |  |  |  |  |
|  |  |  |  |  | C79 | C93 |
| **Total disulfide bonds** |  |  |  |  |  |  |
| **Capsid** | 955<br>(6<sub>C281-C563</sub> x 80 hexons + 5<sub>C97-C271</sub> x 11 pentons + 2<sub>C271-C347</sub> x 210 hexon-hexon interfaces) |  |  |  |  |  |
| **Tail** | 1032<br>(6<sub>C234-C264</sub> x 58 MTP rings + 6<sub>C60-C79</sub> x 57 inter-rings + 6<sub>C131-C19</sub> x 57 inter-rings) |  |  |  |  |  |
| **Tail-TT** | 6 |  |  |  |  |  |
| **Tail-Dit** | 6 |  |  |  |  |  |
| **Dit** | 36<br>(6<sub>C44-C90</sub> + 6<sub>C311-C394</sub> + 6<sub>C322-C337</sub>) |  |  |  |  |  |
| **Dit-β-sandwiches 1-3** | 12<br>(1<sub>C6-C40</sub> x 6 β-sandwich 2 + 1<sub>C12-C305</sub> x 6 β-sandwich 1) |  |  |  |  |  |
| **Virion** | 2047 |  |  |  |  |  |

